# C-terminal evolutionary remodelling of isoleucyl-tRNA synthetases is a prokaryote-specific strategy for tuning aminoacylation rate

**DOI:** 10.64898/2026.02.03.703510

**Authors:** Petra Modrusan, Alojzije Brkic, Andre Lecona Buttelli, Marc Leibundgut, Igor Zivkovic, Nenad Ban, Liam M. Longo, Ita Gruic-Sovulj

## Abstract

Aminoacyl-tRNA synthetases are the guardians of translational fidelity. Their complex function is mirrored by an elaborate structure, which includes multiple nested domains. While the evolutionary pressures that promoted the emergence of some domains, such as the editing domain, are clear, the pressures acting on other domains, particularly those at the C-terminus, are not. Here, we use a combination of kinetic analysis, X-ray crystallography, and bioinformatics to unveil the history and evolutionary forces that have shaped isoleucyl-tRNA synthetase (IleRS) domain structure. We find that the traditional classification into IleRS1 and IleRS2, based on the C-terminal tRNA-recognition domains, is incomplete, as it fails to capture features of the synthetic domain. Guided by the crystal structure of the *Priestia megaterium* IleRS2:tRNA complex, we removed key interactions between IleRS2 and its cognate tRNA and characterised their impact on enzyme activity. We found that D-loop interactions with the IleRS2 C-terminal region are non-essential in prokaryotes, and their loss can even increase catalytic turnover. Further, the zinc-binding domain of IleRS1 recognises the anticodon less stringently than the canonical C-terminal domain of IleRS2. Our data suggest that C-terminal evolutionary remodelling of IleRSs is an ongoing process with a historical precedent, consistent with selection for faster aminoacylation rate.

## Introduction

Aminoacyl-tRNA synthetases (AARSs) are indispensable enzymes that establish the genetic code by coupling amino acids and their cognate tRNAs for ribosomal protein synthesis^1,2^. As guardians of translational accuracy, AARSs are under strong evolutionary pressure for high selectivity in amino acid^3^ and tRNA recognition^4^. In cases where initial amino acid selectivity alone could not achieve the required translation accuracy, AARSs evolved hydrolytic editing^2,5^ to prevent error propagation.

AARSs are divided into two evolutionarily distinct classes, I and II, each comprising ten members^6,7^. Isoleucyl-tRNA synthetase (IleRS) is a class I enzyme responsible for charging tRNA^Ile^ with isoleucine (Ile)^8^ (**Fig. 1A**). IleRS enzymes catalyse an ATP-dependent two-step aminoacylation reaction, in which the amino acid is first activated as an adenylate and then transferred to its cognate tRNA. Both of these steps occur within the synthetic site of the HUP domain^9^ (**Fig. 1B)**. When IleRS fails to reject non-cognate amino acids resembling Ile at the activation step, the enzyme either clears the misactivated amino acids within the HUP domain^10,11^ or hydrolyses the mischarged tRNA^Ile^ at the editing domain (ED)^12,13^ (**Fig. 1A and B**).

**Figure 1.**
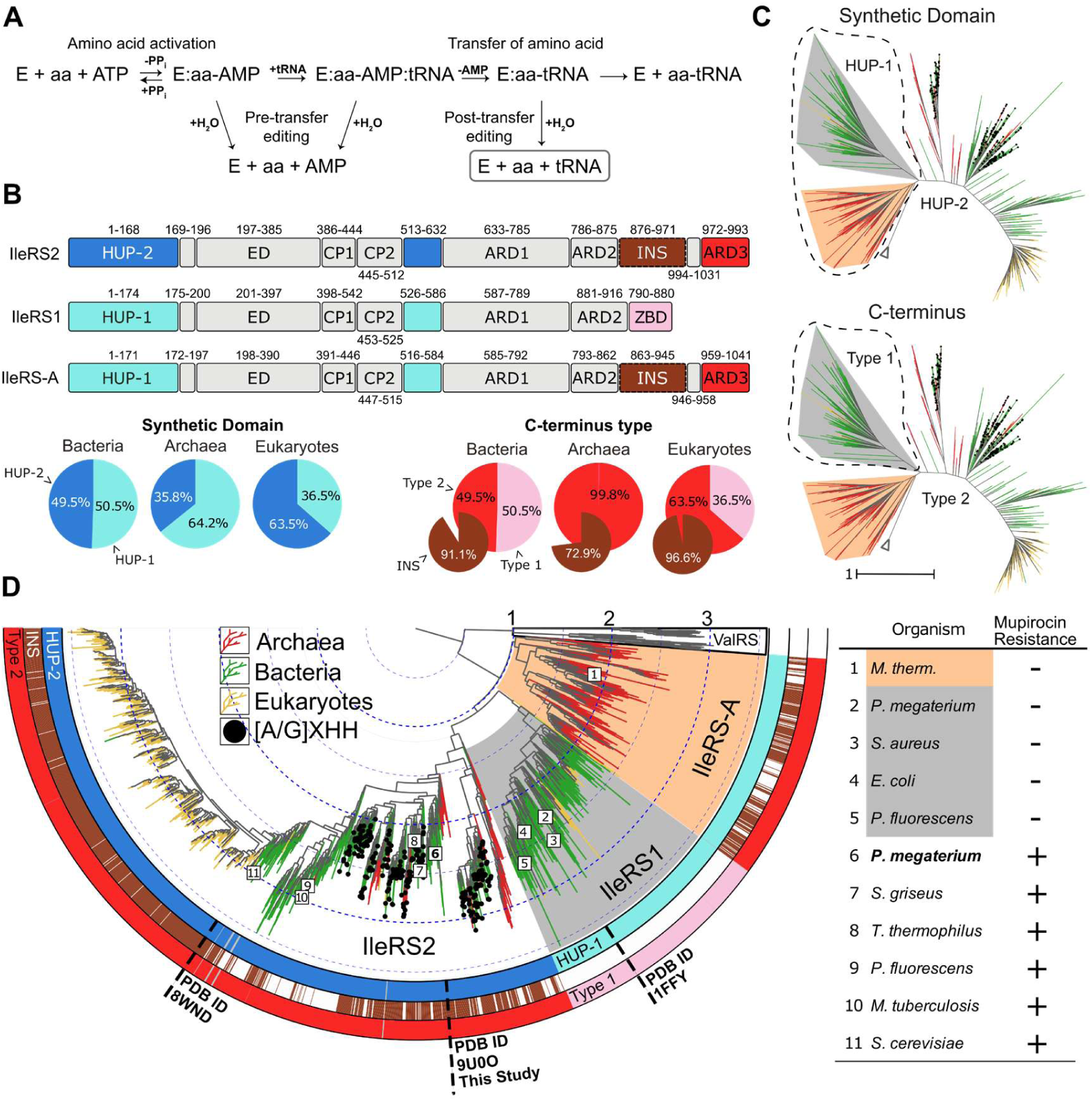
Reaction mechanisms, domain composition, and phylogeny of IleRS. **A)** The IleRS reaction cycle. An amino acid is first activated to form an aminoacyl-adenylate intermediate, followed by aminoacyl transfer to the cognate tRNA within the synthetic site of the HUP domain. If a non-cognate amino acid is activated, it can be cleared within the synthetic site or transferred to tRNA. Mischarged tRNA is rapidly deacylated at the editing domain (ED). **B)** Domain composition for the three primary IleRS groups, which form distinct evolutionary clades. Each group consists of a HUP domain, divided by two connective peptides (CP) and an editing domain (ED), followed by three tRNA anticodon-recognition domains (ARD1–ARD3, ZBD). IleRS-A has a HUP domain that is more closely related to that of IleRS1 (termed HUP-1) but shares a type 2 C-terminal domain organisation with IleRS2, which comprises ARD1-ARD3 and optionally an ∼80 amino acid insertion within the ARD2 domain (INS). In contrast, IleRS1 has a treble clef zinc-binding domain (ZBD) and lacks INS, which is referred to as the type 1 C-terminal organisation. The indicated residue ranges are according to IleRS1 and IleRS2 from *P. megaterium* and IleRS-A from *Methanobacterium thermoautotrophicum*. The phyletic distributions of HUP and C-terminal domain types are presented as pie charts following the same colour scheme as the domains. **C)** Unrooted maximum-likelihood phylogenetic tree of the HUP domain. Branches are coloured according to domains of life (red: archaea; green: bacteria; yellow: eukaryotes). The clades are coloured according to IleRS groups (orange: IleRS-A; grey: IleRS1; white: IleRS2). ValRS sequences are indicated by a collapsed clade (white triangle). In the top and bottom trees, the clades corresponding to the HUP-1 domain type and the type 1 C-terminal architecture are circled for clarity. **D)** Outgroup-rooted maximum-likelihood tree of the IleRS HUP domain. Tree branches and clades are coloured as in (C). Black circles at the leaf tips indicate the presence of the HIGH motif variant [A/G]XHH, which has been shown to confer mupirocin hyper-resistance in IleRS2. From innermost to outermost, the rings annotate the HUP type, the presence of an INS subdomain, and the type of C-terminal organisation, respectively, following the same colour scheme as in (B). IleRS enzymes for which sensitivity to mupirocin has been experimentally assessed^15,20–22,26–30^ are indicated by numbers in the phylogenetic tree, with the corresponding organisms listed in the adjacent table.

IleRSs are traditionally further subdivided into two groups, IleRS1 and IleRS2, based on the distinct anticodon-recognition domains (ARDs) at their C-termini^14–17^, although this classification does not fully reflect their evolutionary relationships (**Fig. 1B**). IleRS1 enzymes have a C-terminal zinc-binding treble-clef motif^18^ (ZBD) and are found only in bacteria and eukaryotic organelles^19^. In contrast, IleRS2 enzymes, present in archaea, the eukaryotic cytoplasm, and many bacteria^15,20,21^, have a C-terminal ARD3 domain that is structurally homologous to the preceding domain (ARD2)^17,22^. IleRS2 can also have a unique insertion in the ARD2 domain, termed INS.

IleRS1 and IleRS2 also differ in their sensitivity to mupirocin, a naturally occurring and clinically relevant polyketide antibiotic^23^ that binds to the synthetic site of the HUP domain^24^. Mupirocin strongly inhibits IleRS1^25^, whereas IleRS2 from eukaryotes^26,27^ and bacteria^15,21,28,29^ display significantly greater mupirocin resistance, requiring >10^3^ higher concentrations to be inactivated. In addition, IleRS2 may harbour a natural, non-canonical variant of the class I AARS signature HIGH motif that confers hyper-resistance to mupirocin^22^.

The fact that IleRS2 in bacteria is a housekeeping enzyme that also confers antibiotic (hyper)resistance raises the question of why IleRS1 has persisted in some bacterial lineages. A clue may be provided by the study of *Priestia megaterium*, one of the rare organisms that has both IleRS1 and IleRS2. Previously, we found that the IleRS1 of *P. megaterium* has a higher aminoacylation turnover rate and supports faster *in vitro* translation than its IleRS2. Moreover, an analysis of reported bacterial doubling times suggested that faster-growing bacteria preferentially employ IleRS1^28^. Could the increase in turnover be related to the presence of ZBD and/or the absence of INS in IleRS1?

Using structural, kinetic, and phylogenetic analyses, we sought to clarify the factors and mechanisms that drove the divergence of IleRS. First, we show that, although IleRS enzymes have traditionally been subdivided into two groups based on their C-terminal domains, they are better described as having split into three evolutionary clades. In addition to bacterial IleRS1 and archaeal/bacterial/eukaryotic IleRS2, we propose a clade of deep-branching archaeal IleRS that we term IleRS-A. Our phylogenetic analysis indicates that IleRS1 diverged from IleRS-A, likely driven by a replacement of the C-terminal domain. We also find evidence for ongoing C-terminal remodelling, with a concerted loss of the INS subdomain across the prokaryotic tree of life.

To better understand the role of ARD3 and INS, and the forces shaping their evolution, we solved the crystal structure of *P. megaterium* IleRS2 in complex with tRNA and performed a detailed kinetic characterisation of ARD3, INS, and tRNA mutants. We find that *P. megaterium* IleRS2 recognises the anticodon bases with slightly higher fidelity than *P. megaterium* IleRS1. We also show that INS, which is present exclusively in the ARD2 domain of IleRS2 and IleRS-A, first served as a tRNA-binding domain in archaea and bacteria but later evolved into a tRNA recognition element in eukaryotes. Finally, we re-trace the evolutionary loss of the INS subdomain, showing that it is a relatively facile evolutionary transition. In summary, our data frame the replacement of ARD3 by ZBD to form the IleRS1 clade and the ongoing loss of INS as prokaryotic strategies for increased aminoacylation turnover.

## Results

### Resolving a missing deep-branching archaeal IleRS clade

The traditional subdivision of the IleRSs based on differences in their C-terminal domains has generally been interpreted as reflecting an evolutionary split into two clades: IleRS1 found in bacteria versus IleRS2 found in archaea, some bacteria, and the cytoplasm of eukaryotes^19^. However, deeper phylogenetic analysis revealed that some archaeal IleRSs, so far attributed to the IleRS2 clade, actually cluster more closely with IleRS1^21,22^. This observation raises the question of whether C-terminal domain types correspond to a clade separation.

To clarify the evolutionary history of IleRS, we constructed a database of 96,219 IleRS sequences spanning bacteria, archaea, and eukaryotes and annotated their protein domain architectures (see **Methods** and **Fig. S1** for more details). Phylogenetic trees were calculated for each of the core structural domains (HUP, ED, CP1, CP2, and ARD1; **Fig. 1B**) and rooted with ValRS as the outgroup. For each structural domain, representative sequences were re-sampled with replacement five times, aligned, and phylogenetic trees were inferred (see **Methods** for more details). Unrooted and outgroup-rooted trees of the IleRS synthetic (HUP) domain are presented in **Fig. 1C** and **D**, respectively, while trees calculated using other structural domains are presented in **Fig. S2.**

Foremost, we find that the standard two-group division scheme does not conform to the clade structure of the phylogenetic tree. In addition to IleRS1 and IleRS2, we identify a novel clade of archaeal IleRS (IleRS-A). This clade has a C-terminal domain organisation like IleRS2 but has a HUP domain that is more closely related to that of IleRS1 (HUP-1; **Fig. 1C-D**). Indeed, the closer resemblance of IleRS-A to IleRS1 is consistent across phylogenetic trees inferred from the HUP domain, as well as trees calculated from other core domains (**Fig. S2**). The one IleRS-A tested to date was found to be sensitive to mupirocin^20^, consistent with IleRS1, further supporting the classification of IleRS-A as having a HUP-1 synthetic domain. Additionally, the non-canonical HIGH motif characteristic for IleRS2 was neither detected in IleRS-A nor in IleRS1.

For nearly all trees from all protein domains (**Fig. 1** and **Fig. S2**), the ValRS-outgroup-inferred root is positioned closest to, or within, IleRS-A. Although the position of IleRS1 with respect to IleRS-A genes is not entirely stable, all trees suggest that IleRS1 emerged from an IleRS-A-like ancestor. Remodelling of the C-terminus likely occurred concomitant with, or in close temporal proximity to, the emergence of IleRS1, likely in a bacterial ancestor. Eukaryotic IleRS1 (in organelles) and IleRS2 genes then evolved independently from bacterial ancestors. Thus, current evidence supports a model in which IleRS evolution involved two independent passages from archaeal to bacterial to eukaryotic forms.

### C-terminal domain exchange and loss have shaped, and are shaping, IleRS evolution

The phylogenetic tree shows that remodelling of the C-terminus is a core driver of IleRS diversification. Two key examples emerge: exchange of ARD3 for a ZBD with an unrelated fold and loss of the INS subdomain (∼80 residues). As the ZBD domain is somewhat shorter and less complex than the ARD3 domain (31 vs. 82 residues on average, respectively), both of these processes result in a modest shortening of IleRS. However, whereas ARD3 exchange appears to have occurred just once, relatively early in the evolution of contemporary IleRS proteins, INS subdomain loss appears to be an active, ongoing process: we find that a number of organisms have lost the INS subdomain independently in both the IleRS-A and IleRS2 clades, with the IleRS2 eukaryote sub-clade being a notable exception (**Fig. 1D**).

Why are the C-terminal domains of IleRS a hotspot for diversification, and what forces drove, and are still driving, these changes? To address these questions, we sought to (a) identify the residues of ARD3 and INS domains that interact with tRNA; (b) evaluate the catalytic consequences of these interactions; and (c) experimentally realise an event of C-terminal remodelling.

### The tRNA^Ile^ recognition by bacterial IleRS2

Because eukaryotic INS subdomains are rarely lost, we reasoned that the role of the INS domain in eukaryotes may differ from that in bacteria or archaea. Thus, we solved the crystal structure of the IleRS2 from *P. megaterium* (*Pm*IleRS2) bound to *Escherichia coli* tRNA^Ile^ and ATP (**Fig. 2A, Table S1**). This is the first crystal structure of an IleRS that contains the more evolutionarily unstable INS domain, making it a particularly useful point of comparison with the tRNA-bound structure of yeast IleRS2 ^17^ and the tRNA- and mupirocin-bound structure of *Staphylococcus aureus* IleRS1 (*Sa*IleRS1)^16^.

**Figure 2.**
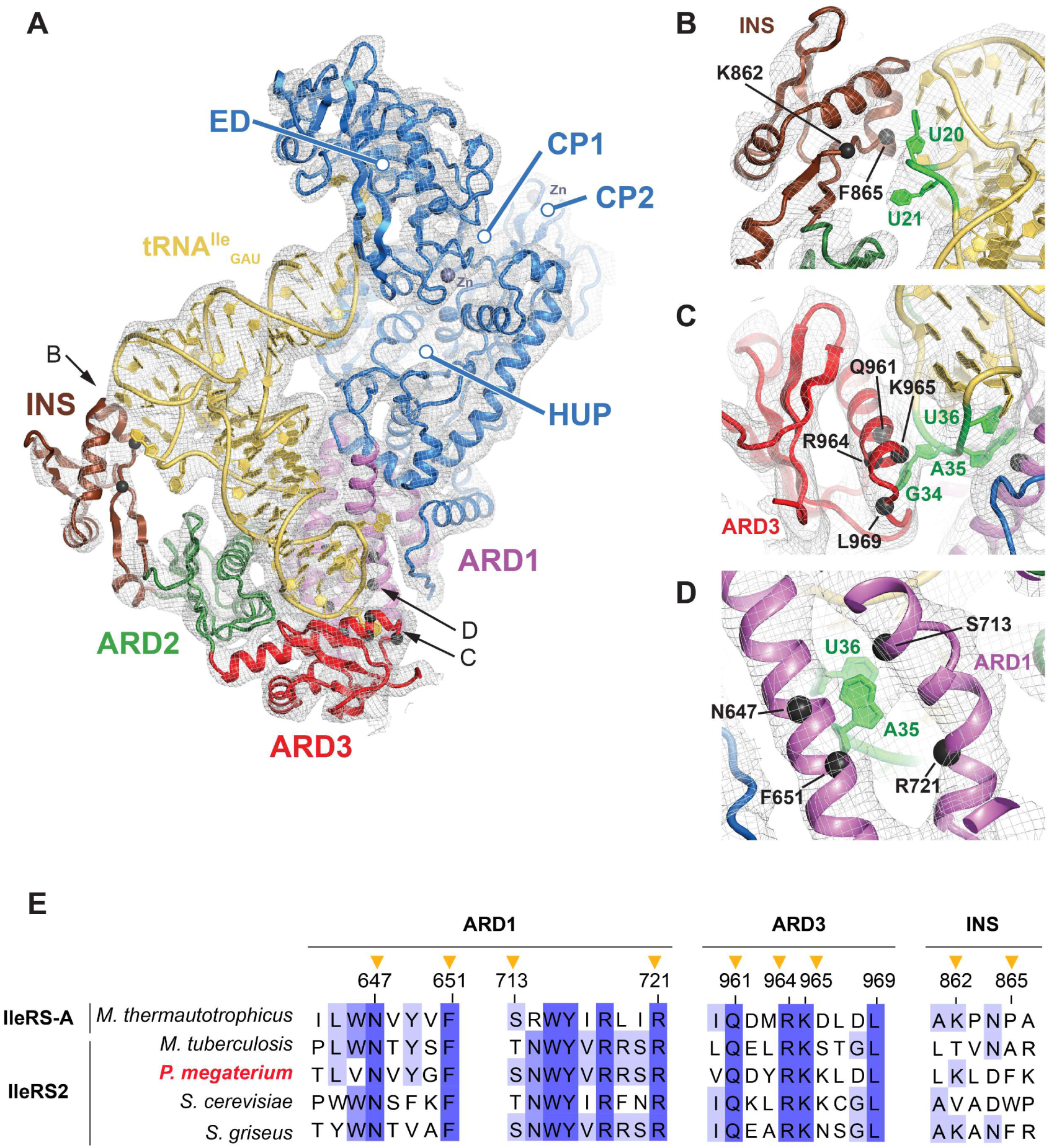
tRNA recognition by *Pm*IleRS2. **A)** Structure of the *Pm*IleRS2:*Ec*tRNA^Ile^:ATP complex. As the precise interactions between the side chains of amino acid residues and the tRNA were not resolved, the participation of the residues (depicted as spheres for their C*α*) in tRNA recognition was instead inferred from comparison with the related crystal structure from yeast (PDB: 8WND; **Fig. S6**) and corroborated by detailed kinetic analysis (**Table 1**). *Pm*IleRS2 is colored according to domain borders, and the 2m*F*_o_–D*F*_c_ Fourier difference map (grey mesh) is carved 4.6 Å around the model and contoured at 0.9 *σ* (panels A,C and D) or 0.6 *σ* (panel B). **B-D)** D-loop and anticodon recognition by *Pm*IleRS2. Insertion into ARD2 (INS) establishes interactions with the D-loop, likely with U20 and U21 nucleotides. (B). The ARD3 likely recognises the first anticodon base, G34, via the pocket comprising residues R964, L969, Q961, and K965 (C), while the second and third bases, A35 and U36, are recognised by R721, N647, F651, and S731 at the ARD1 subdomain (D). **E**) Conservation of the *Pm*IleRS2 residues putatively involved in tRNA recognition. The sequence alignment shows that the residues involved in anticodon recognition are highly conserved, whereas those involved in D-loop recognition are less conserved. Mutated residues are marked with arrows.

**Table 1.**
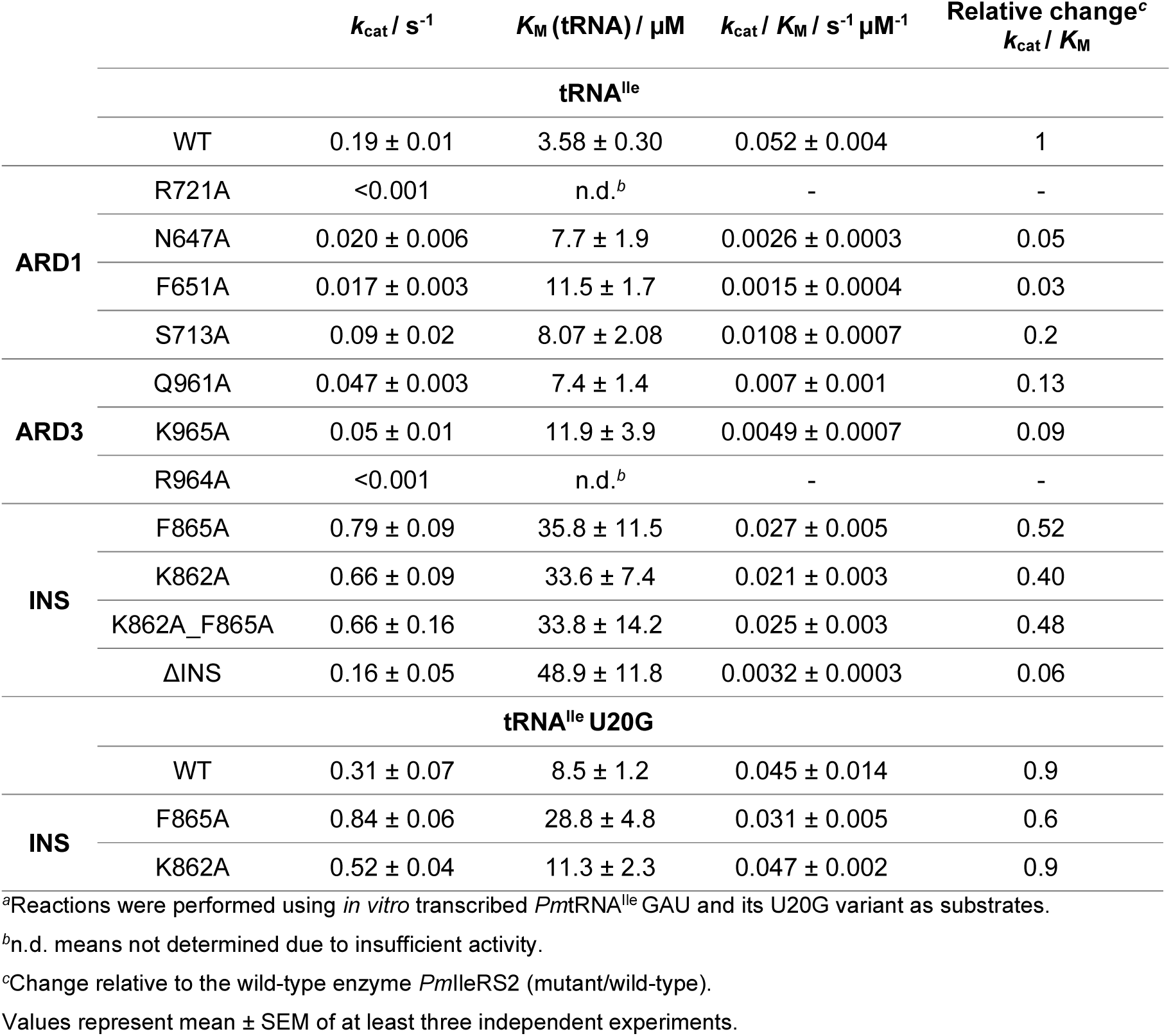
Steady-state kinetic parameters for aminoacylation of wild-type and D-loop tRNA^Ile^ variants by wild-type *Pm*IleRS2 and its mutants.^a^.

Despite a limited resolution of 5.3 Å (**Table S1**), all domains of *Pm*IleRS2, as well as the bound tRNA, were visible in the electron density of all four complexes in the asymmetric unit (**Fig. S3**). This allowed modelling of the protein and tRNA backbones using the structures of *Pm*IleRS2:Ile-AMS (PDB: 8C8V)^22^ and yeast IleRS2 bound to the tRNA^Ile^ and Ile (PDB: 8WND)^17^ as guides. After iterative readjustment and restrained refinement using non-crystallography symmetry (NCS) averaging, the final model displayed high-quality metrics at the achieved resolution (**Table S1**), supporting its accuracy and robustness. Further, the structural model was functionally verified by extensive mutagenesis and kinetic measurements (detailed below).

In the *Pm*IleRS2:tRNA complex (**Fig. 2A**), the tRNA acceptor stem is captured in the editing site, and the anticodon loop is lodged within a cleft formed by the ARD1-ARD3 domains. The overall architecture is consistent with previous tRNA-bound IleRS structures^16,17^ (**Fig. S4**) and with a type 2 C-terminal domain organisation (**Fig. 1B**). When compared to the structure of *Pm*IleRS2 without tRNA (PDB: 8C8V)^22^ we find that tRNA binding promotes a ∼77° rotation of the INS subdomain of all four complexes in the asymmetric unit (**Fig. S5**), enabling its engagement with the D-loop. Our model suggests that K862 and F865 (both in the INS subdomain) are positioned close to the bases of U20 and U21 and their connecting phosphate backbone (**Fig. 2B**).

To recognise the anticodon loop, our model indicates that ARD3 (**Fig. 2A**) may read the first anticodon base, G34, through the pocket formed by the conserved residues R964, L969, and Q961 (**Fig. 2C**), as observed in yeast (**Fig. S6**). Likewise, the second and third anticodon nucleotides, A35 and U36, are likely recognised by ARD1 residues R721, S713, F651 and N647 (**Fig. 2D**).

To corroborate the structural data, we investigated the role of residues putatively involved in tRNA recognition using mutagenesis and steady-state kinetic analysis. Substitution of R964, possibly engaged in G34 recognition (**Fig. 2C**), abolished the capacity of IleRS2 to charge tRNA^Ile^ with Ile, confirming its key role in G34 recognition. The aminoacylation rate for the R964A mutant was below 0.001 s^-1^, demonstrating a 10^3^-fold drop relative to the wild-type (WT) (**Table 1**). The barely detectable activity of the R964A mutant precluded the determination of a *K*_M_ (tRNA). We next mutated two nearby, highly conserved ARD3 residues: Q961, which is likely involved in G34 recognition, and K965, which is thought to stabilise Q961. The catalytic efficiency (*k*_cat_/*K*_M_) for the Q961A and K965A mutants decreased 10-fold relative to WT (**Table 1**), suggesting a supporting role in G34 recognition by R964.

The next two anticodon bases, A35 and U36, may be recognised by the strictly conserved R721 within the ARD1 domain of *Pm*IleRS2 (**Fig. 2D**). Similar to R964, the R721A mutation decreased the aminoacylation rate to below 0.001 s^-1^ (**Table 1**). Mutation of the ARD1 residues N647, F651, and S713 to alanine (**Fig. 2D**) decreased the aminoacylation *k*_cat_/*K*_M_ by 5 to 30-fold, mainly via a *k*_cat_ effect (**Table 1**). These results demonstrate that R721 is critical for A35 and U36 recognition, with other residues in ARD1 further refining *Pm*IleRS2 specificity.

### Anticodon recognition by *Pm*IleRS2 is more stringent than by *Pm*IleRS1

A comparison of tRNA recognition between *Sa*IleRS1^16^ and *Pm*IleRS2 (**Fig. 2A**) revealed distinct tRNA binding modes, in which *Pm*IleRS2 interacts with the D-loop, while *Sa*IleRS1 engages the D-stem of the tRNA (**Fig. S4**). In contrast, binding modes are highly similar for yeast^17^ and bacterial *Pm*IleRS2 (**Fig. S4**). Although IleRS1 and IleRS2 both utilise ARD1 and ARD3 to read the anticodon triplet, *Sa*IleRS1 recognises the G34 base via backbone and stacking interactions (**Fig. S6**), whereas *Pm*IleRS2 likely uses specific side-chain recognition, as observed in yeast (**Fig. S6**). Contacts with A35 involve analogous residues from ARD1 in both IleRSs, whereas the reading of U36 differs between them. Protein contacts to U36 present in *Pm*IleRS2 and yeast IleRS2 are replaced in *Sa*IleRS1 by interactions with U33 of the tRNA itself (**Fig. S6**). Thus, the structural data indicate that *Pm*IleRS2 exhibits stricter tRNA anticodon recognition than *Sa*IleRS1.

To explore this hypothesis by kinetics, we produced anticodon variants of *P. megaterium* tRNA^Ile^ GAU and tested whether they could be aminoacylated by both IleRS1 and IleRS2 from *P. megaterium* (**Fig. 3**, **Table S2**). As expected, the tRNA variants exhibit a significant reduction in steady-state aminoacylation by *Pm*IleRS1 (also shown for *E. coli* IleRS1^31^) as well as *Pm*IleRS2, demonstrating the key role of the anticodon in tRNA recognition by both enzymes. Interestingly, the tRNA variants were aminoacylated 3-15-fold less efficiently by *Pm*IleRS2 (**Fig. 3 inset**), indicating that IleRS2 exhibits more stringent anticodon recognition, in line with the structural data. The smallest difference between the two IleRSs was observed for the A35U variant, consistent with analogous reading, whereas G34C, U36A, and U36C variants show larger differences of similar magnitude.

**Figure 3.**
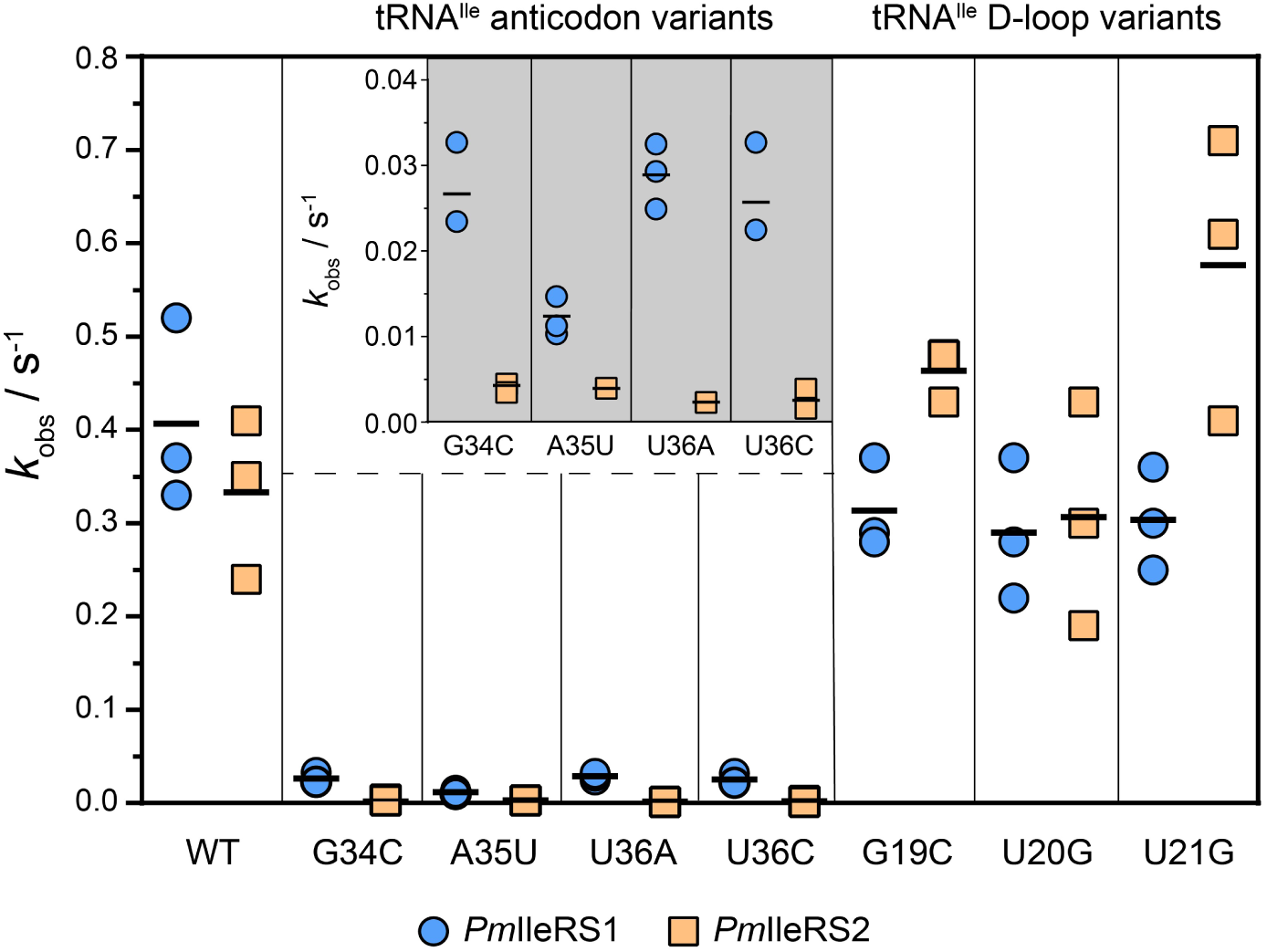
Comparison of the *Pm*tRNA^Ile^ anticodon and D-loop variants in aminoacylation by *Pm*IleRS1 (blue circles) and *Pm*IleRS2 (orange rectangles). The *k*_obs_ is obtained at a saturating concentration of tRNA^Ile^ (50 μM), thus providing a close approximation of *k*_cat_. The inset shows an enlarged view of the tRNA anticodon variants. Mean values of at least three independent experiments are indicated with a dash; the standard error of the mean is given in **Table S3**. tRNA U20G was chosen as a representative for detailed kinetic analysis (**Table 1**).

### Anticodon recognition is critical for the isoleucyl transfer step

Although the R721A and R964A IleRS2 mutants cannot form Ile-tRNA^Ile^ (**Table 1**), their *k*_cat_/*K*_M_ values for Ile activation, which is the first step in Ile-tRNA^Ile^ formation, remain comparable to the WT enzyme (**Table S3**), demonstrating that the mutations influence only the second step, aminoacyl transfer. The transfer of the aminoacyl moiety to tRNA could be assessed under single-turnover conditions by mixing the preformed IleRS:Ile-AMP complex with a limiting amount of tRNA^Ile^. The IleRS:Ile-AMP complex was formed *in situ* by incubating IleRS with Ile and ATP before mixing with the tRNA.

Single-turnover analysis revealed that R964A and R721A mutants both exhibit a 10^4^-fold drop in isoleucyl transfer rate relative to the WT enzyme (**Fig. 4A**). Mutating the tRNA instead of the enzyme returned a similar outcome: the WT enzyme exhibited a 10^3^-fold lower rate of aminoacyl transfer to a tRNAU36C anticodon variant than to WT tRNA (**Fig. 4A**). Thus, anticodon recognition is critical for assembling a catalytically competent conformation for the transfer step, consistent with earlier findings that tRNA identity elements preferentially stabilise the transition state for the transfer step^32,33^.

**Figure 4.**
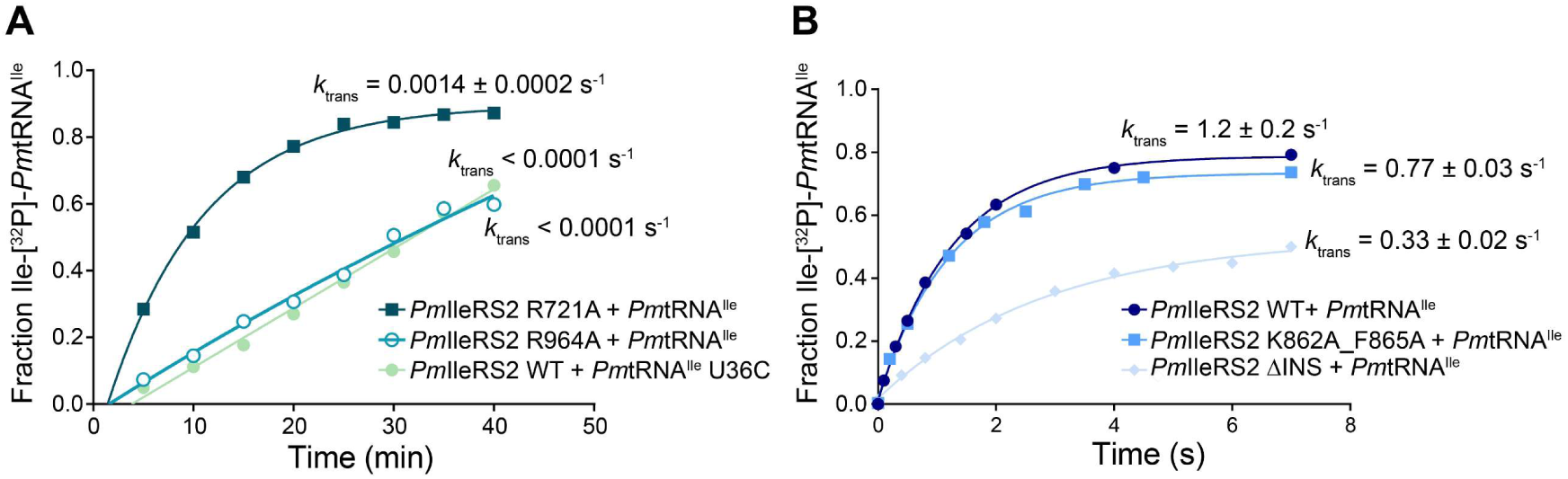
Single-turnover analysis of aminoacyl transfer mutants of *Pm*IleRS2 and tRNA^Ile^. **A)** Mutation of the anticodon base U36 and the anticodon-reading residues R721 and R964 gave a 10^3^- or 10^4^-fold slower transfer step, respectively. Doubling the enzyme concentration returned the same rate, showing that chemistry, not binding, is followed (data not shown). **B)** Variations in INS yield up to a 5-fold decrease in the transfer step. Time points on a minute time scale (A) were obtained by manual mixing, while the ones on millisecond and second time scales (B) were sampled using a rapid chemical quench (RQF) instrument. Values represent mean ± SEM of at least three independent experiments.

### The INS subdomain mediates base-non-specific contacts with the tRNA^Ile^ D-loop

To explore interactions of D-loop nucleotides U20 and U21 with the INS subdomain of *Pm*IleRS2 (**Fig. 2B**), we characterised several U20/U21 tRNA^Ile^ variants, including U20G, U20C, U21G, and U21C. We also characterised the G19C variant, as the D-loop conformation might be disturbed by the absence of the G19:C56 Watson-Crick base pair. The tRNA variants showed no decrease in the aminoacylation rate constants relative to the WT tRNA (**Fig. 2**, **Table S2**), indicating that the D-loop establishes base-nonspecific binding interactions with the INS subdomain, a phenomenon known as an indirect readout^34^. Interestingly, the D-loop variants generally showed a slight *increase* in the rate of aminoacylation by *Pm*IleRS2 relative to WT tRNA (**Fig. 3**, **Table S2**). A detailed steady-state analysis using tRNA U20G revealed increases in both *k*_cat_ and *K*_M_ (**Table 1**). An increase in *k*_cat_ is unexpected for tRNA variants, suggesting that the D-loop does not function as a tRNA identity element, and thus, unlike the anticodon, it is not essential for productive juxtaposition of catalytic moieties in the transition state of the aminoacyl transfer step (see below).

No change in turnover of the D-loop variants was observed when they were charged by *Pm*IleRS1 (**Fig. 3**, **Table S2**). This result was expected, as IleRS1 does not establish contacts with the tRNA D-loop. Thus, the small but consistent increase in *k*_cat_ by *Pm*IleRS2 is not related to some change in tRNA structure but is specific to tRNA recognition by IleRS2.

### INS is a tRNA-binding element but not a recognition determinant in *Pm*IleRS2

The intriguing finding that D-loop variants show *increased k*_cat_ (**Table 1**) prompted us to explore the effect of amino acid substitutions in the INS subdomain of *Pm*IleRS2, which should affect binding analogously. To this end, we characterised the aminoacylation activities of the K862A and F865A single mutants and the K862A_F865A double mutant. All mutants exhibit a ∼4-fold *increase in k*_cat,_ accompanied by a 10-fold rise in *K*_M_ (**Table 1**), mirroring the effect observed with the tRNA U20G variant (**Table 1**). Thus, loss of D-loop interactions increases *k*_cat_, suggesting that these interactions restrain a rate-limiting step in the *Pm*IleRS2 catalytic cycle. Earlier findings demonstrated that Ile-tRNA^Ile^ dissociation limits IleRS catalytic turnover^35^, indicating that impairment of the D-loop interactions may enhance Ile-tRNA^Ile^ dissociation.

To test the requirement for an intact INS and attempt to realise the ongoing INS loss process inferred from the phylogenetic tree (**Fig. 1D**), we produced a mutant in which the entire INS subdomain (residues 851-925) was replaced with a short (GGGGS)_2_ linker (ΔINS). The ΔINS protein variant did not exhibit a noticeable difference in its overall fold relative to WT, as assessed by circular dichroism (**Fig. S7**), suggesting that INS makes only a modest contribution to maintaining the IleRS2/IleRS-A architecture. This result is consistent with ΔINS exhibiting only a small reduction in *k*_cat_ and no *K*_M_ effect for Ile in the activation step (**Table S3**). In contrast, truncation increases the *K*_M_ for tRNA in aminoacylation by 15-fold (**Table 1**), consistent with INS participating in D-loop binding (**Fig. 2**).

The second step of aminoacylation (the isoleucyl transfer step) can be impaired when recognition of the tRNA identity elements is disrupted^32^. Yet, both the K862A_F865A and ΔINS mutants display only a 2- to 5-fold decrease in the single-turnover rate constants relative to the WT enzyme (**Fig. 4B**). This confirms that, in contrast to anticodon recognition (**Fig. 4A**), *Pm*IleRS2 interactions with the D-loop are not essential to form the catalytically competent conformation for aminoacyl transfer. Instead, the INS subdomain primarily enhances tRNA affinity.

### INS is not mandatory for the functioning of *Pm*IleRS2 *in vivo*

To explore whether INS is required *in vivo*, we assessed the capacity of INS mutants to rescue *E. coli* growth in the presence of mupirocin, an inhibitor of its sole endogenous IleRS1. The K862A, F865A, and K862A_F865A mutants complement *E. coli* growth to a similar extent as WT *Pm*IleRS2 (**Fig. S8A),** even at low IPTG concentration, such as 2 μM (**Fig. 5A**). Complementation by the ΔINS mutant was more challenging (**Fig. S8B**) and requires a 100-fold higher IPTG concentration as well as co-overexpression of *Pm*tRNA^Ile^ in the cell (**Fig. 5B**). This coincides with the *in vitro* data showing that ΔINS exhibits lower *k*_cat_/*K*_M_ compared to other INS mutants due to the lack of a positive effect on *k*_cat_. Thus, our data provide evidence that interactions between INS and the tRNA D-loop are non-essential under the tested conditions, and that even the entire INS subdomain is not mandatory for *Pm*IleRS2 functioning *in vivo* when expressed at sufficient levels.

**Figure 5.**
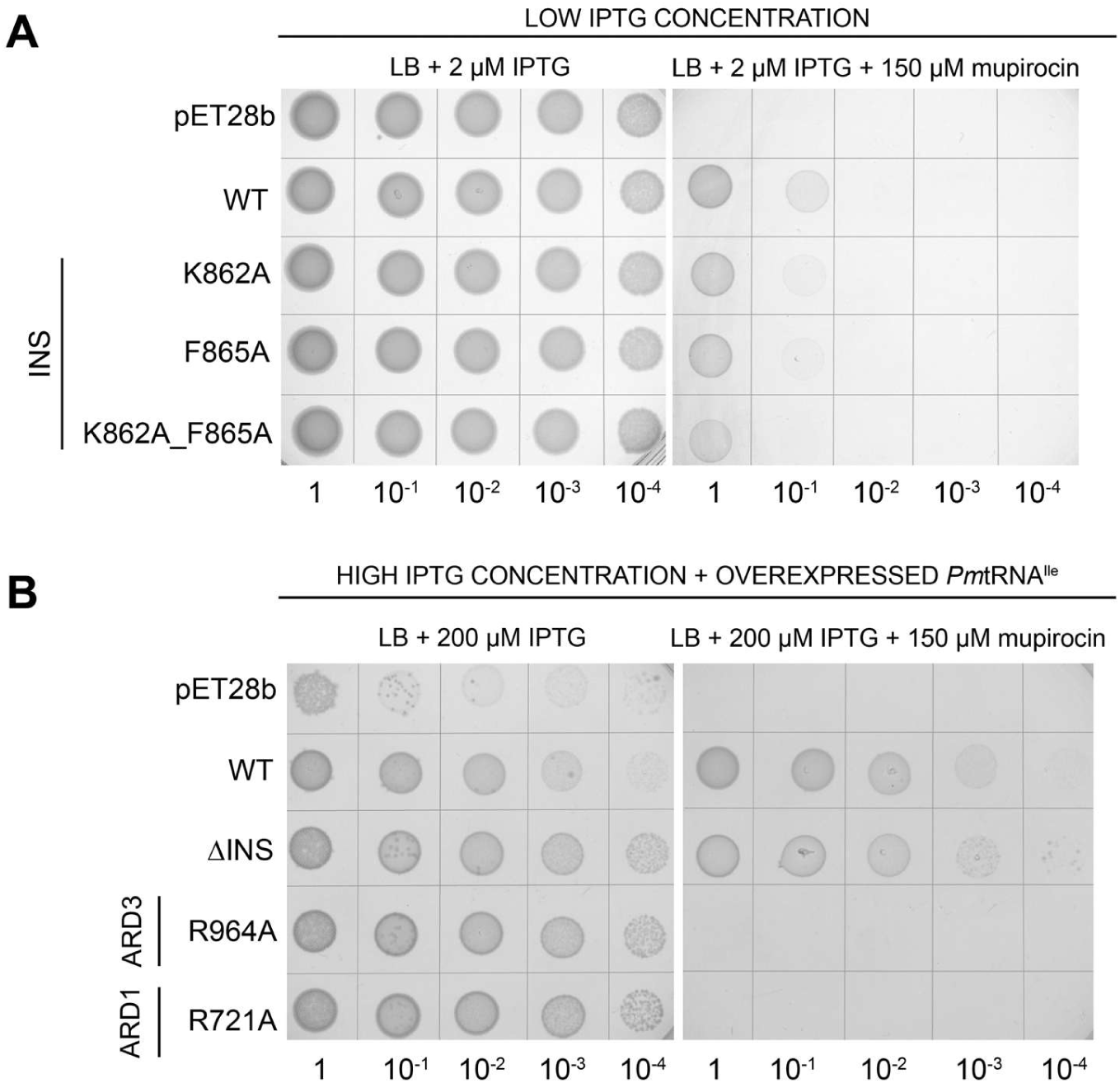
Complementation of *E. coli* growth by *Pm*IleRS2 variants in the presence of mupirocin. Functionality of the *Pm*IleRS2 mutants *in vivo* was assessed by their ability to complement mupirocin-inhibited *E. coli* IleRS1. Expression of the mutants was induced from the pET28b plasmid using various concentrations of IPTG with or without co-overexpression of *Pm*tRNA^Ile^ (**Fig. S8**). **A-B**) INS mutants rescue *E. coli* growth already at a 2 μM IPTG (A), while ΔINS requires 200 μM IPTG and co-overexpression of tRNA^Ile^ (B). *Pm*IleRS2 anticodon mutants are unable to complement *Ec*IleRS1, even at 200 μM IPTG and with co-overexpression of tRNA^Ile^ (B). For each panel, growth without mupirocin is shown on the left, the negative control (empty plasmid with and without overexpressed tRNA^Ile^) is positioned at the top, and the positive control (overexpression of the WT) is placed in the second row. The experiments were performed in at least two independent replicates.

In sharp contrast, ARD1-based R721A and ARD3-based R964A mutants could not complement the activity of mupirocin-inhibited *E. coli* IleRS1, even at high IPTG induction (**Fig. S8B**) and tRNA co-overexpression (**Fig. 5C**), in agreement with their abolished anticodon recognition *in vitro* (**Table 1**). Finally, the remaining IleRS2 mutants complement *E. coli* (**Fig. S8C**) in line with *in vitro* aminoacylation efficiencies (**Table 1**).

## Discussion

### IleRS split into three clades based on the HUP domain and C-terminus

IleRSs have traditionally been subdivided into two groups based on their C-terminal domains, primarily based on the presence of a Zn-binding domain (ZBD) that is exclusive to IleRS1 (**Fig. 1B**). IleRS1 (historically referred to as bacterial IleRS) is found in bacteria and eukaryotic organelles, whereas IleRS2 (historically referred to as archaeal/eukaryotic IleRS) resides in archaea, the cytoplasm of eukaryotes, and bacteria, though initial reports suggested modest bacterial occurrence.^19^ Later studies revealed that a substantial number of bacteria encode IleRS2^21^. Although the subdivision into two groups was thought to reflect phylogenetic clades, we observed that some archaeal IleRS, classified as IleRS2 by their C-terminal domain, are more closely related to IleRS1^21,22^. This result indicated that the C-terminal domain is not the only driver of IleRS phylogeny.

To address the issue more thoroughly, we conducted a comprehensive phylogenetic analysis and inferred trees based on the different structural modules of IleRS (**Fig. S2**). The HUP domain-based tree, along with other trees, identified three distinct IleRS clades (**Fig. 1C and D**). Two of the clades align with the traditional classification: i) IleRS1 enzymes present in bacteria and eukaryotic organelles, and ii) IleRS2 enzymes found in archaea, bacteria, and eukaryotic cytoplasm. However, a novel third clade, designated here as IleRS-A, comprises archaeal enzymes that possess a type 2 C-terminal domain (i.e., lacking a ZBD) but have an HUP domain that groups more closely with that of IleRS1 (referred to as HUP-1). Two defining features of HUP-1 are sensitivity to the antibiotic mupirocin^20,24^ and near-absolute conservation of the catalytic HIGH motif. In contrast, HUP-2, assigned to the IleRS2 enzymes, exhibits mupirocin resistance, which can be further augmented with a natural variation of the HIGH motif, [A/G]IHH^22^. Presented data suggest that the split to HUP-1 and HUP-2 predates the emergence of mupirocin, which seems to originate in *Pseudomonas*^23^, with the HUP-2 (IleRS2) lineage likely being naturally predisposed to develop resistance.

Our comprehensive analysis offers a novel view of IleRS phylogeny, showing that the early archaeal split gave rise to the IleRS-A and IleRS2 lineages, followed by the subsequent emergence of IleRS1 within the IleRS-A branch. The deep-rooted position of IleRS-A strongly suggests that its HUP-1 domain more closely resembles the ancestral IleRS synthetic domain. Thus, the type 2 C-terminal domain architecture likely evolved first, presumably through duplication of ARD2 to generate ARD3. The emergence of IleRS1, then, coincided with the acquisition of ZBD and the loss of ARD3. The observation that all IleRS1 enzymes lack an INS subdomain indicates that the ancestor of IleRS1 resembled an IleRS-A-like enzyme lacking the insertion into ARD2.

### The C-terminal domain reorganisation in IleRS1 was likely driven by demands for faster aminoacylation

Differentiating between the AUA codon for Ile and the AUG codon for methionine (Met) is challenging, because the uridine at the wobble (34) position of the tRNA^Ile^ cannot discriminate between A and G under standard wobble pairing rules^36^. To uncouple decoding of AUA from AUG, different evolutionary lineages employ distinct post-transcriptional modifications at position 34 of tRNA^Ile^ ^37–39^. Aside from rare cases where tRNA^Ile^ UAU is present, and the ribosome resolves AUA versus AUG codon ambiguity on its own^40^, bacteria encode two isoacceptors. The major isoacceptor, tRNA^Ile^ GAU, decodes AUU and AUC, while the minor, tRNA^Ile^ CAU, would pair with AUG if left unmodified. Accurate Ile incorporation is ensured by two mechanisms. First and foremost, C34 of the tRNA^Ile^ CAU isoacceptor is post-transcriptionally modified to lysidine (L), which redirects its pairing specificity to the AUA codon and acts as a strong positive determinant for recognition by IleRS. Second, the unmodified tRNA^Ile^ CAU is not recognised by IleRS but is instead charged with Met by MetRS^41^. Because this mischarged Met-tRNA^Ile^ CAU reads the AUG codon, it does not compromise translational fidelity.

Recognition of the wobble position during aminocylation is mediated by the C-terminal domain of IleRS, which differs between IleRS1 and IleRS2/IleRS-A. It has been proposed^17^ that the ZBD domain was acquired to recognise a bacteria-specific lysidine modification^42,43^. However, the authors overlooked that nearly half of the bacteria harbour IleRS2 instead of IleRS1^21^ (**Fig. 1B)**, and utilise ARD3 for tRNA recognition, suggesting that ZBD in bacteria must serve a different purpose.

What other factors might have driven the acquisition of the ZBD in IleRS1? Aminoacylation of the tRNA anticodon variants reveals that *Pm*IleRS1 is less sensitive to the base identity than *Pm*IleRS2 (**Fig. 3**). This difference may reflect a trade-off between accuracy and velocity in IleRS1, a principle well-documented in translation fidelity^44–47^. Could the absence of INS in IleRS1 (**Fig. 1B**) stem from similar evolutionary pressures? Bacterial IleRS2 mutants with impaired D-loop interactions display an unexpected increase in *k*_cat_ (**Table 1**), suggesting that the lack of INS in IleRS1 might also signify a shift towards faster product release, which is the rate-limiting step in IleRS turnover^35^. Therefore, the selection of ZBD and loss of INS could represent an adaptation consistent with increased catalytic velocity of IleRS1 in faster-growing bacteria^28^.

### Transition of the INS subdomain from a tRNA binder to a catalytic mediator

The stages of INS emergence could not be resolved unambiguously (**Fig. 1D**). However, because we found no evidence for an INS-like insertion in ARD3, INS most likely emerged after the ARD2 duplication event that gave rise to ARD3, but prior to the occurrence of the last common ancestor of IleRS. Indeed, many of the deepest-branching IleRS-A genes contain INS, suggesting that the last common ancestor of IleRS likely possessed this insertion. Given that diverse bacteria and archaea now lack the INS subdomain, we conclude that contraction of IleRS through INS loss is an actively ongoing process in prokaryotes. Taken together, these data suggest that INS loss likely reflects a reversion to an IleRS domain organisation that predates the last common IleRS ancestor.

Our experimental data could explain why the INS subdomain in bacteria is evolutionarily unstable, while it becomes more established in eukaryotes. In *Pm*IleRS2, mutations of INS residues F865 and K862 increase the *K*_M_ for tRNA (**Table 1**), but only lead to a modest reduction in the rate of the aminoacyl transfer step (**Fig. 4B**). This indicates that, while INS interactions with the D-loop enhance tRNA binding affinity, they do not facilitate maturation of the catalytically productive active site conformation. Because the D-loop is not an identity element in bacteria and likely also archaea, some IleRS2 and IleRS-A enzymes, may naturally lose INS (**Fig. 1B**), while in those that retain it, like *Pm*IleRS2, INS is not essential for activity either *in vitro* (**Table 1**) or *in vivo* (**Fig. 5B**). Note that because the aminoacyl transfer step is not rate-limiting, the INS mutants (F865A and K862A) actually display *increased k*_cat_ (**Table 1**). Along this line, remarkably, deletion of the entire subdomain results in an enzyme (ΔINS) with an unchanged turnover number (**Table 1**), providing a rationale for how INS may undergo dynamic loss in prokaryotes.

In contrast to bacteria, the D-loop in yeast apparently serves as an identity element of tRNA^Ile^. Yeast IleRS2 specifically recognises the U20 base via K915 (**Fig. S6**), and removing this interaction using D-loop variants of the tRNA^Ile^ significantly decreases the aminoacylation rate.^17^ In line with this evolutionary shift, INS loss events are notably less common in eukaryotes than in bacteria and archaea (**Fig. 1B**). Thus, we propose that INS initially emerged as a tRNA-binding module with a minor role in juxtaposing reaction moieties for efficient catalysis. Later, in eukaryotes, INS appears to have evolved to play a significant role in adopting the catalytic conformation required for the aminoacyl transfer step, thereby becoming essential.

## Materials and methods

Protein sequences from COG0060 (IleRS) were obtained from the EggNOG database v5.0^48^. Sequences were clustered at 60% sequence identity using CD-HIT v4.8.1^49^ [cd-hit -c 0.6 -n 4]. Cluster representatives were aligned with MAFFT L-INS-i v7.505^50^ [mafft --localpair --maxiterate 1000 --thread -1] and trimmed using TrimAl v1.4.rev15^51^ [trimal -gappyout]. The alignment was visually inspected with JalView v2.11.5.0^52^ and outliers were removed manually. Finally, sequences from previously crystallised IleRS1 (PDB 8C9E) and IleRS2 (PDB 8C8V)^22^ were added to the alignment [mafft --add], for a total of 2,212 representative IleRS sequences. A phylogenetic tree was inferred for the representative IleRS sequences using IQ-TREE v2.4.0^53^ Model Finder Pro (MFP)^54^, and ultra-fast bootstrap (UFBoot)^55^ [iqtree -m MFP -B 1000 -T 1] using a single core^56^. Based on the phylogenetic tree, the sequences were divided into two groups, corresponding to IleRS2 and IleRS-A (1047 sequences) and IleRS1 (488 sequences), and re-aligned. Using previously reported domain boundaries^22^, reference crystal structures, secondary structure propensities, and conservation scores, the alignments were segmented into the canonical IleRS domains: HUP, connective peptide 1 (CP1), CP2, editing domain (ED), and anticodon-recognition domain 1 (ARD1). Alignments for the C-terminal domains of IleRS1 (IleRS1_ARD2, and IleRS1_ZBD) and IleRS2/IleRS-A (IleRS2/IleRS-A_ARD2, IleRS2/IleRS-A_ARD3 and IleRS2/IleRS-A_INS) were also generated. Each domain alignment was used to construct a hidden Markov model (HMM) using the HMMER suite v3.3.2^57^ [hmmbuild]. Sequences from the closely related AARS families ValRS (COG0525), LeuRS (COG0495), and MetRS (COG0143) were obtained from the EggNOG database and clustered as above. IleRS HMM profiles were then analysed by searching [hmmsearch --domtblout] against ValRS, LeuRS, and MetRS to assess the extent of discrimination at the domain level (**Fig. S1**).

### IleRS sequence database construction

The IleRS domain HMM profiles were used to search the Genome Taxonomy Database (GTDB) v220.0^58^ and the EukProt v3^59^ database. Using a cutoff of 0.75 profile coverage and 0.01 iE-value, domain annotations were assigned with preference given to non-overlapping hits with lower iE-values. The domain annotation script, DOTATE v1.2.0, is available on GitHub (https://github.com/AndreLecona/Dotate). Selection criteria for candidate IleRS sequences were (a) matching at least four of the five canonical IleRS core domains (HUP, CP1, ED, CP2, and ARD1) and (b) matching at least one IleRS type-specific domain (IleRS1_ARD2, IleRS2/IleRS-A_ARD2, IleRS1_ZBD, IleRS2/IleRS-A_ARD3, IleRS2/IleRS-A_INS). In total, 7,360 archaeal, 109,435 bacterial, and 1,588 eukaryotic proteins were classified as potential IleRSs.

Once again, COG IleRS, ValRS, LeuRS, and MetRS reference sequences were clustered as above and added to a custom sequence database using BLAST v2.17.0^60^ [makeblastdb]. Candidate IleRS sequences were clustered at 70%, 60%, and 50% sequence identity for eukaryotic, archaeal, and bacterial genes, respectively, using CD-HIT as above. Representative genes were searched against the labelled COG sequences [blastp -outfmt "6 qseqid sseqid pident length evalue bitscore qlen slen" -evalue 1e-5] and a sequence similarity network (SSN) was constructed by filtering hits with pident < 30 and coverage < 0.5. Network visualizations were performed in Cytoscape v3.10.4^61^ using the perfuse force directed layout weighting by bit-score. Only sequences whose representative clustered with a COG IleRS were retained (**Fig. S1**), resulting in a total 97,676 IleRS sequences.

Filtered IleRS sequences were classified according to their C-terminal domain configurations and compiled into a validated IleRS database integrating data from *Bacteria*, *Archaea* and *Eukarya*. From this database, domain composition analyses were calculated.

### Phylogenetic analysis

Representative sequences from *Archaea* and *Bacteria* in the IleRS database were randomly selected at the family level according to the GTDB taxonomy. Bacterial representatives were clustered at 60% sequence identity as above. The final dataset included all eukaryotic IleRS sequences, family-level representatives for *Archaea*, and family-level clustered representatives for *Bacteria*, providing an approximately balanced representation across the three domains of life. ValRS sequences from COG were clustered and aligned as above, and the homologous domains for HUP, CP1, ED, CP2, and ARD1 were manually extracted. 50 ValRS sequences, composed of 25 bacterial and 25 archaeal representatives, were included as outgroups for tree rooting. Multiple sequence alignments were generated for IleRS domain sequences, then merged using MAFFT –merge [mafft –localpair –maxiterate 1000 --merge] with the corresponding ValRS alignment. Then the merged alignment was trimmed as above using TrimAl -automated1 option (optimised for Maximum Likelihood phylogenetic tree calculation). Phylogenies were then inferred using IQ-TREE as above. To ensure the robustness of the phylogenetic inferences, the representative sampling procedure was repeated for each new domain to be analysed (*i.e.*, family representatives were re-randomised, re-clustered, re-aligned, and new trees were calculated). The phylogeny for each domain was calculated five times, with -MFP only applied in the first run. The resulting model for that domain was reused across the other four runs (HUP: LG+R10, CP1: LG+R9, ED: LG+R9, CP2: LG+R9, ARD1: LG+G4). Consistency of topological features across these replicates, as well as across domains, was used as a measure of robustness. For each domain, all five consensus trees were topologically consistent, and the trees with the highest log-likelihood are presented in **Fig. S2**.

### Preparation and purification of biological material for crystallisation

Preparation of *Pm*IleRS2 suitable for crystallisation experiments is described elsewhere^22^. *Escherichia coli* tRNA^Ile^ GAU (*Ec*tRNA^Ile^ GAU) was transcribed in *E. coli* BL21(DE3) for 15 h at 30 °C using 1 mM IPTG from a ΔpET3a-derived plasmid containing the tRNA gene. The first base pair of the *Ec*tRNA^Ile^ GAU was changed from A–U to G–C to improve transcription yield, without affecting kinetic competence^12^. LB was supplemented with 5 mM MgCl₂ to improve the yield. The crude tRNA was purified through sequential phenol:chloroform extraction, polyethylene glycol precipitation, deacylation, and dialysis, according to established protocols. Isolation of high-homogeneity tRNA^Ile^ GAU from the crude tRNA preparation was conducted by reversed-phase chromatography as previously described^10,11^. Fractions with 95% or higher tRNA^Ile^ purity were pooled, dialysed against 5 mM HEPES-KOH (pH 7.5 at 20 °C), lyophilised, and stored at −80 °C. The final preparation for crystallisation had an isoacceptor content exceeding 98% as determined by aminoacylation.

### Crystallization

Wild-type *Pm*IleRS2 was diluted to 19.77 mg/mL in 25 mM HEPES-KOH (pH 7.5, 20 °C), 50 mM NaCl, and 10 mM β-mercaptoethanol. *Ec*tRNA^Ile^ GAU stock (99.34 mg/mL in 5 mM HEPES-KOH, pH 7.5, 20 °C) was heat-renatured by dilution with MgCl₂ to 55.1 mg/mL tRNA and 100 mM MgCl₂. The ternary complex was assembled by mixing 38.5 µL *Pm*IleRS2, 6.5 µL renatured tRNA, and 5 µL 100 mM ATP (in protein buffer), yielding a ∼1:2 protein:tRNA molar ratio with 10 mM ATP. Crystals were grown at 20°C by sitting-drop vapour diffusion, using 1+1 µl drops of complex and reservoir solutions (50 mM HEPES-NaOH, pH 6-8, 50 mM MgCl_2_, 7.5–15% PEG3350, 0.1 mM NH_4_OAc). After 2-7 days, the crystals were stabilised by adding 25% pentaerythritol propoxylate (5/4 PO/OH) in one step. After incubation for 5 minutes, the crystals were mounted on Hampton loops and flash-frozen in liquid nitrogen.

### Data collection and processing

Diffraction data were collected at beamline X06SA (Swiss Light Source, Paul Scherrer Institut, Villigen, Switzerland) using a Dectris Eiger 16M detector and a wavelength of 1.0 Å. A total of 3600 frames (0.1° oscillation angle) were indexed and scaled with XDS and XSCALE^62^ into space group P2_1_2_1_2_1_ (unit cell: 121.75 × 147.01 × 344.11 Å; α = β = γ = 90°), yielding usable data to 5.3 Å resolution based on CC₁_/_₂ criterion^63^ (**Table S1**).

### Phasing and refinement

An initial molecular replacement solution was obtained using PHASER^64^ with a search model assembled from the *Pm*IleRS2:Ile-AMS structure (PDB: 8C8V)^22^ and including the tRNA from the superimposed *Sa*IleRS1:tRNA:mupirocin structure (PDB: 1FFY)^16^, revealing four copies in the asymmetric unit (ASU) with a solvent content of ∼59%. Clear m*F*_o_–D*F*_c_ Fourier difference density was observed for omitted tRNA and protein parts after rigid body refinement in PHENIX^65^ using the individual *Pm*IleRS2 domains and tRNAs as groups (**Fig. S3A**). The quality of the map was improved by a four-fold real-space averaging in CCP4^66^, which allowed completion of the tRNA model (**Fig. S3B**). The less-well-resolved domains of the protein were repositioned manually by rigid body docking into unbiased OMIT maps and rebuilding of the connections. The model was further improved by adjusting peripheral loops according to m*F*_o_–D*F*_c_ difference Fourier maps. The improvement of the phases was monitored by calculation of anomalous difference Fourier maps for the zinc ions in the zinc fingers (**Fig. S3C** and **S3D**), which resulted in increasing peak sigma levels with decreasing R_work_ and R_free_ values, and by calculation of simulated annealing composite OMIT maps to minimise model bias. Because only weak residual m*F*_o_–D*F*_c_ Fourier difference density was observed for the bound ATP cofactors, the ligand was not modelled. Great care was taken to keep the model geometry intact and to prevent over-refinement by only editing the original model and refining it to convergence. This approach allowed progressive improvement of the model-to-map fit with a concomitant decrease in *R*_free_ and *R*_work_ values. To remove side chain clashes between neighbouring copies in the ASU, the completed input model was first subjected to *phenix.geometry_minimization* for 3 macrocycles (each comprising 100 iterations), followed by several cycles of coordinate refinement to convergence with *phenix.refine*, which included torsion-angle NCS and secondary structure restraints as well as the group-wise B factor setting (two groups per residue) and choosing a fixed value of *wxc* = 0.6 for the weighting between the geometry and X-ray targets to prevent overfitting. To take into account the anisotropic motion of domains, the B-factor refinement also including domain-wise TLS, which further improved the R_work_ and R_free_ values by ∼ 1% to 20.5% and 25%, respectively (**Table S1**). The final model, validated using *MolProbity*^67^, exhibited excellent statistics for the resolution (**Table S1**). For figures, the best-ordered complex (chains *A* for protein and *a* for tRNA, respectively) was used.

Comparison of the final model with the recently published 2.8 Å crystal structure of the yeast IleRS2:tRNA^Ile^:isoleucine complex (PDB: 8WND)^17^ revealed only minor differences for the protein, essentially confirming our low-resolution model. However, as some base orientations in the anticodon loop and the acceptor stem of the tRNA described the electron density more accurately, they were consequently readjusted and the structure was re-refined. The deposited final model (PDB accession code 9U0O, corresponding to extended pdb_00009U0O) includes all side chains and is meant to represent a pseudoatomic interpretation of the *Pm*IleRS2:*Ec*tRNA^Ile^:ATP complex.

### Cloning and mutagenesis

The expression vectors for *Pm*IleRS2 and *Pm*tRNA^Ile^ GAU mutants were generated by whole-plasmid amplification of pET28b(+)_*Pm*IleRS2 or ΔpET3a_*Pm*tRNA^Ile^ GAU using *Phusion High-Fidelity DNA Polymerase* (Thermo Scientific) according to the manufacturer’s recommendations. Mutations were introduced by site-directed mutagenesis using custom-designed primers (**Table S4**). All mutants were verified by sequencing (Macrogen inc.).

### Production and purification of proteins

Wild-type *Pm*IleRS2 and its mutant variants were overexpressed in *E. coli* BL21(DE3) as described elsewhere^22^. The cells were harvested by centrifugation and resuspended in 10 or 25 mL of washing buffer (20 mM Hepes-KOH, pH = 7.5, 500 mM NaCl, 20 mM MgCl_2_, 10 mM imidazole, 10 mM 2-mercaptoethanol), depending on the method used for cell lysis. Two methods were employed for cell lysis: sonication^22^ and high-pressure homogenization (Constant Systems Cell Disruptor BT40/T52/AA) at 20,000 psi (1378 bar). The soluble protein fraction was loaded onto a HisTrapHP column (Cytiva, 1 or 5 mL) and the protein was eluted with 200 mM imidazole. A further purification step included size-exclusion chromatography on a Superdex Increase 200 10/300 GL (Cytiva) or HiLoad Superdex 200 16/600 (Cytiva) column in a buffer containing 20 mM Hepes-KOH (pH 7.5), 50 mM NaCl, 10% glycerol, and 10 mM 2-mercaptoethanol. Fractions with monomeric protein were pooled, concentrated to ∼20 mg/mL and stored at -80 °C. Proteins were expressed from two or three distinct clones, purified and tested independently in each kinetic assay.

### *In vitro* transcription and purification of tRNA^Ile^

The DNA template required for *in vitro* transcription was amplified by PCR using the plasmid ΔpET3a_*Pm*tRNA^Ile^ GAU, *Phusion* DNA polymerase, an upstream primer: 5’-TCTCAAGGGCATCGGTCGAC-3’, and a specific 2’-*O*-methylated downstream primer: 5’-TmGGTGGGCCTAAGTGGACTC-3’. *In vitro* transcription was performed at 37 °C for 4 hours in a 1 mL reaction mixture containing 1.8 μg of DNA template, 40 mM Tris-HCl (pH 8.0), 20 mM MgCl_2_, 2 mM spermidine, 10 mM DTT, 0.005 mg/mL BSA, 5 mM NTPs, 0.008 U/μL TiPP, 0.05 U/μL RNase, and 0.1 mg/mL T7 RNA polymerase. The resulting tRNA transcript was purified by preparative denaturing gel electrophoresis on a 6% (w/v) polyacrylamide gel. After electrophoresis, tRNA was extracted from the gel using a buffer containing 50 mM Tris-HCl (pH 7.5), 10 mM MgCl_2_, 1 mM EDTA, and 0.1% SDS. The transcript was then precipitated with 96% ice-cold ethanol and 3 M NaOAc (pH 5.2), followed by centrifugation and washing with 70% ice-cold ethanol. Finally, tRNA was dialyzed against 10 mM Hepes-KOH (pH 7.5) and kept at -20 °C. Labelled [^32^P]-tRNAs were prepared as described elsewhere^13^. Transcripts were renatured before use by heating at 85 °C for 3 minutes, followed by slow cooling to reaction temperature.

### Amino acid activation assay

Amino acid activation assay was monitored by an ATP-PP_i_ exchange assay using radioactively labelled γ-[^32^P]ATP^68^. The reactions were carried out at 30 °C in a buffer containing 20 mM Hepes-KOH (pH 7.5), 10 mM MgCl_2_, 0.1 mg/mL BSA, 5 mM DTT, 4 mM ATP, and 1 mM PP_i_. The enzyme was present at 20 or 100 nM, and Ile was used at concentrations ranging from 0.1 to 10 times the *K*_M_ values. The reaction was stopped at various time points with 0.5 M NaOAc (pH 5.2) and 0.1% SDS, followed by product separation by TLC and quantification as described^28^. The *k*_cat_ and *K*_M_ (Ile) were obtained by fitting the data to the Michaelis–Menten equation using GraphPad Prism 6.

### Aminoacylation assay

Aminoacylation assay was performed at 30 °C in a reaction mixture containing 20 mM Hepes-KOH (pH 7.5), 150 mM NH_4_Cl, 10 mM MgCl_2_, 0.008 U/μL TIPP, 0.01 mg/mL BSA, 5 mM DTT, and 2 mM ATP. The enzyme was present at concentrations ranging from 10 to 2000 nM. The steady-state reaction was started by adding 4 mM Ile to mixtures containing tRNA^Ile^ at concentrations ranging from 0.1 to 10 times the *K*_M_. The reaction was stopped at various time points by 0.1 M NaOAc and 5% HOAc, and the quenched aliquots were treated with P1 nuclease^69^, followed by product separation by TLC and quantification as described^28^. The *k*_cat_ and *K*_M_ (tRNA) were obtained by fitting the data to the Michaelis–Menten equation using GraphPad Prism 6. For tRNA^Ile^ anticodon and D-loop variants, reactions were carried out at a saturating concentration of tRNA^Ile^ (50 μM) and 40 μM [^14^C]Ile (50-100 mCi/mmol). At the indicated time points, aliquots were spotted on Whatman 3MM filter discs, quenched in ice-cold 10% TCA, and washed twice with cold 5% TCA. The amount of [^14^C]Ile-tRNA^Ile^ retained on the filters was quantified by liquid scintillation counting. The reaction rate was determined from a linear plot of product concentration versus time.

### Single-turnover transfer of an amino acid to tRNA

The transfer of isoleucine to tRNA^Ile^ was measured by mixing IleRS:aa-AMP complex preformed *in situ* with a limiting concentration of [^32^P]tRNA^Ile^. 20-40 μM enzyme was first incubated for 10 minutes at 30 °C with 4 mM Ile and 4 mM ATP in a buffer containing 20 mM Hepes-KOH (pH 7.5), 150 mM NH_4_Cl, 10 mM MgCl_2_, 0.008 U/μL TIPP, 0.01 mg/mL BSA, and 5 mM DTT to allow for IleRS:aa-AMP complex formation. The transfer reaction was initiated by mixing the solution with an equal volume of 2-4 μM [^32^P]tRNA^Ile^ and stopped at various time points with 0.1 M NaOAc and 5% HOAc. The quenched aliquots were treated with P1 nuclease, followed by product separation by TLC and quantisation as described^69^. The *k*_trans_ was extracted by fitting the experimental data into a single exponential equation *y* = *Y*_O_ + *A* x (1 − *e*^-ktrans x t^) using GraphPad Prism 6.

### Complementation assay

Overnight cultures were prepared from *E. coli* BL21(DE3) cells transformed with a single plasmid expressing the enzyme of interest or co-transformed with both the enzyme-expression plasmid and the *Pm*tRNA^Ile^-expression plasmid. The overnight cultures were diluted to an OD_600_ of 1 for the complementation assay. The cultures were then subjected to five serial dilutions. Aliquots of 10 µL from each dilution were spotted onto LB plates, which were supplemented with 30 µg mL^-1^ kanamycin, 100 µg mL^-1^ ampicillin, 1 mM MgCl_2_, and 0.1 mM ZnCl_2_ with or without the addition of 150 µM mupirocin. IPTG was added to LB agar medium at concentrations ranging from 2 µM to 200 µM. The plates were grown for 24 h at 30 °C.

## Authors contribution

IGS, LML, and NB conceived and supervised the project. PM produced WT and mutant enzymes and tRNAs, performed kinetic analysis and complementation assays with guidance from IZ. AB produced material for structural analysis and solved the crystal structure together with ML. ALB performed the phylogenetic analysis. IGS and LML wrote the manuscript with contributions from all authors.

## Data availability

The structural data generated in this study have been deposited in the Protein Data Bank (PDB) under accession number: 9U0O (*Pm*IleRS2:*Ec*tRNA^Ile^ GAU:ATP).

## Acknowledgements

We are grateful to Bartol Božić for the construction of ΔINS and to Julia Jelena Toplak for her assistance in purifying materials for crystallography. We are indebted to Jeff Errington, Morana Dulić, and Marko Močibob for their comments on the manuscript.

## Funding

This work was supported by the Croatian Science Foundation under the project number HRZZ-IP-2022-10-1400 (IGS), the Swiss National Science Foundation under the project number 310030E_215868 (NB), the European Regional Development Fund (infrastructural project CIuK) [KK.01.1.1.02.0016], and the Human Frontier Science Program Grant Number RGEC29/2025 (LML). LML and IGS acknowledge travel support from ELSI for collaborative exchange.

## Competing interests

The authors reveal no competing interests.

## Materials & Correspondence

Correspondence and material requests should be addressed to Ita Gruic-Sovulj, Liam M. Longo, or Nenad Ban.

## Notes

### Competing Interest Statement

The authors have declared no competing interest.

### Summary of Updates

Figure legends (Figures 3, 4, and 5) updated to include additional experimental information; Materials and Methods section updated with protein production details.

## References

1. Angel, M., Gomez, R. & Ibba, M. Aminoacyl-tRNA synthetases. RNA 26, 910–936 (2020).

2. Perona, J. J. & Gruic-Sovulj, I. Synthetic and editing mechanisms of aminoacyl-tRNA synthetases. Top. Curr. Chem. 344, 1–41 (2014).

3. Tawfik, D. S. & Gruic-Sovulj, I. How evolution shapes enzyme selectivity – lessons from aminoacyl-tRNA synthetases and other amino acid utilizing enzymes. FEBS Journal 287, 1284–1305 (2020).

4. Giege, R. & Eriani, G. The tRNA identity landscape for aminoacylation and beyond. Nucleic Acids Res. 51, 1528–1570 (2023).

5. Zivkovic, I., Dulic, M. & Gruic-Sovulj, I. Mechanisms and kinetic assays of aminoacyl-tRNA synthetases. FEBS Lett. (2025).

6. Eriani, G., Delarue, M., Poch, O., Gangloff, J. & Moras, D. Partition of tRNA synthetases into two classes based on mutually exclusive sets of sequence motifs. Nature 347, (1990).

7. Ribas de Pouplana, L. & Schimmel, P. Two Classes of tRNA Synthetases. Cell 104, 191–193 (2001).

8. Nordin, B. E. & Schimmel, P. Isoleucyl-tRNA Synthetases. in The Aminoacyl-tRNA Synthetases (eds. Ibba, M., Francklyn, C. & Cusack, S.) 24–35 (2005).

9. Gruic-Sovulj, I., Longo, L. M., Jabłońska, J. & Tawfik, D. S. The evolutionary history of the HUP domain. Crit. Rev. Biochem. Mol. Biol. 57, 1–15 (2022).

10. Dulic, M., Cvetesic, N., Perona, J. J. & Gruic-Sovulj, I. Partitioning of tRNA-dependent editing between pre- and post-transfer pathways in class I aminoacyl-tRNA synthetases. Journal of Biological Chemistry 285, 23799–23809 (2010).

11. Dulic, M., Perona, J. J. & Gruic-Sovulj, I. Determinants for tRNA-dependent pretransfer editing in the synthetic site of isoleucyl-tRNA synthetase. Biochemistry 53, 6189–6198 (2014).

12. Cvetesic, N., Bilus, M. & Gruic-Sovulj, I. The tRNA A76 hydroxyl groups control partitioning of the tRNA-dependent pre- and post-transfer editing pathways in class I tRNA synthetase. Journal of Biological Chemistry 290, 13981–13991 (2015).

13. Zivkovic, I., Ivkovic, K., Cvetesic, N., Marsavelski, A. & Gruic-Sovulj, I. Negative catalysis by the editing domain of class I aminoacyl-tRNA synthetases. Nucleic Acids Res. 50, 4029–4041 (2022).

14. Landro, J. A. & Schimmel, P. Zinc-dependent cell growth conferred by mutant tRNA synthetase. Journal of Biological Chemistry 269, 20217–20220 (1994).

15. Sassanfar, M., Kranz, J. E., Gallant, P., Schimmel, P. & Shiba, K. A Eubacterial Mycobacterium tuberculosis tRNA Synthetase Is Eukaryote-like and Resistant to a Eubacterial-Specific Antisynthetase Drug. Biochemistry 35, (1996).

16. Silvian, L. F., Wang, J. & S, T. A. Insights into Editing from an Ile-tRNA Synthetase Structure with tRNAIle and Mupirocin. Science (1979). 285, (1999).

17. Chen, B. et al. The mechanism of discriminative aminoacylation by isoleucyl-tRNA synthetase based on wobble nucleotide recognition. Nature Communications 15, (2024).

18. Krishna, S. S., Majumdar, I. & Grishin, N. V. Structural classification of zinc fingers. Nucleic Acids Res. 31, 532–550 (2003).

19. Woese, C. R., Olsen, G. J., Ibba, M., Söll, D. & Söll, S. Aminoacyl-tRNA Synthetases, the Genetic Code, and the Evolutionary Process. MMBR 64, 202–236 (2000).

20. Jenal, U. et al. Isoleucyl-tRNA synthetase of Methanobacterium thermoautotrophicum Marburg: Cloning of the gene, nucleotide sequence, and localization of a base change conferring resistance to pseudomonic acid. Journal of Biological Chemistry 266, 10570–10577 (1991).

21. Cvetesic, N. et al. Naturally occurring Isoleucyl-tRNA synthetase without tRNA-dependent pre-transfer editing. Journal of Biological Chemistry 291, 8618–8631 (2016).

22. Brkic, A. et al. Antibiotic hyper-resistance in a class I aminoacyl-tRNA synthetase with altered active site signature motif. Nat. Commun. 14, (2023).

23. Thomas, C. M., Hothersall, J., Willis, C. L. & Simpson, T. J. Resistance to and synthesis of the antibiotic mupirocin. Nat. Rev. Microbiol. 8, 281–289 (2010).

24. Zivkovic, I. & Gruic-Sovulj, I. Exploring mechanisms of mupirocin resistance and hyper-resistance. Biochem. Soc. Trans. 52, 1109–1120 (2024).

25. Hughes, J. & Mellows, G. Inhibition of Isoleucyl-Transfer Ribonucleic Acid Synthetase in Escherichia coli by Pseudomonic Acid. Biochem. J 176, 305–318 (1978).

26. Hughes, J. & Mellows, G. Interaction of pseudomonic acid A with Escherichia coli B isoleucyl-tRNA synthetase. Biochem. J 191, 209–219 (1980).

27. Racher, K. I., Kalmar, G. B. & Borgford, T. J. Expression and characterization of a recombinant yeast isoleucyl-tRNA synthetase. Journal of Biological Chemistry 266, 17158–17164 (1991).

28. Zanki, V., Bozic, B., Mocibob, M., Ban, N. & Gruic-Sovulj, I. A pair of isoleucyl-tRNA synthetases in Bacilli fulfills complementary roles to keep fast translation and provide antibiotic resistance. Protein Science 31, (2022).

29. Yanagisawa, T. & Kawakami, M. How does Pseudomonas fluorescens avoid suicide from its antibiotic pseudomonic acid? Evidence for two evolutionarily distinct isoleucyl-tRNA synthetases conferring self-defense. Journal of Biological Chemistry 278, 25887–25894 (2003).

30. Pope, A. J. et al. Characterization of isoleucyl-tRNA synthetase from Staphylococcus aureus: II. Mechanism of inhibition by reaction intermediate and pseudomonic acid analogues studied using transient and steady-state kinetics. Journal of Biological Chemistry 273, 31691–31701 (1999).

31. Nureki, O. et al. Molecular recognition of the identity-determinant set of isoleucine transfer RNA from Escherichia coli. J. Mol. Biol. 236, 710–724 (1994).

32. Ibba, M., Sever, S., Praetorius-Ibba, M. & Söll, D. Transfer RNA identity contributes to transition state stabilization during aminoacyl-tRNA synthesis. Nucleic Acids Res. 27, 3631–3637 (1999).

33. Uter, N. T., Perona, J. J. & Doudna, J. A. Long-range intramolecular signaling in a tRNA synthetase complex revealed by pre-steady-state kinetics. PNAS 101, 14396–14401 (2004).

34. Perona, J. J. & Hou, Y. M. Indirect readout of tRNA for aminoacylation. Biochemistry 46, 10419–10432 (2007).

35. Eldredt, E. W. & Schimmelt, P. R. Investigation of the Transfer of Amino Acid from a Transfer Ribonucleic Acid Synthetase-Aminoacyl Adenylate Complex to Transfer Ribonucleic Acid. Biochemistry 11, (1968).

36. Crick, F. H. C. Codon—anticodon pairing: The wobble hypothesis. J. Mol. Biol. 19, 548–555 (1966).

37. Nishikawa, K. et al. The Modified Wobble Base Inosine in Yeast tRNAIle Is a Positive Determinant for Aminoacylation by Isoleucyl-tRNA Synthetase. Nature 336, 179–181 (1988).

38. Suzuki, T. et al. Agmatine-conjugated cytidine in a tRNA anticodon is essential for AUA decoding in archaea. Nat. Chem. Biol. 6, 277–282 (2010).

39. Muramatsu, T. et al. A novel lysine-substituted nucleoside in the first position of the anticodon of minor isoleucine tRNA from Escherichia coli. Journal of Biological Chemistry 263, 9261–9267 (1988).

40. Taniguchi, T. et al. Decoding system for the AUA codon by tRNAIle with the UAU anticodon in Mycoplasma mobile. Nucleic Acids Res. 41, 2621–2631 (2013).

41. Muramatsu, T. et al. Codon and amino-acid specificities of a transfer RNA are both converted by a single post-transcriptional modification. Nature 336, (1988).

42. Suzuki, T. & Miyauchi, K. Discovery and characterization of tRNAIle lysidine synthetase (TilS). FEBS Lett. 584, 272–277 (2010).

43. Muraski, M. J. et al. The bacterial tRNA-modifying enzyme tRNAIle lysidine synthetase is genetically conserved but catalytically variable. Journal of Biological Chemistry 301, (2025).

44. Liu, C. et al. Kinetic Quality Control of Anticodon Recognition by A Eukaryotic Aminoacyl-tRNA Synthetase. J. Mol. Biol. 367, 1063–1078 (2007).

45. Wohlgemuth, I., Pohl, C., Mittelstaet, J., Konevega, A. L. & Rodnina, M. V. Evolutionary optimization of speed and accuracy of decoding on the ribosome. Philosophical Transactions of the Royal Society B: Biological Sciences 366, 2979–2986 (2011).

46. Banerjee, K., Kolomeisky, A. B. & Igoshin, O. A. Elucidating interplay of speed and accuracy in biological error correction. Proc. Natl. Acad. Sci. U. S. A. 114, 5183–5188 (2017).

47. Yu, Q., Mallory, J. D., Kolomeisky, A. B., Ling, J. & Igoshin, O. A. Trade-Offs between Speed, Accuracy, and Dissipation in tRNAIle Aminoacylation. Journal of Physical Chemistry Letters 11, 4001–4007 (2020).

48. Huerta-Cepas, J. et al. EggNOG 5.0: A hierarchical, functionally and phylogenetically annotated orthology resource based on 5090 organisms and 2502 viruses. Nucleic Acids Res. 47, D309–D314 (2019).

49. Li, W. & Godzik, A. Cd-hit: A fast program for clustering and comparing large sets of protein or nucleotide sequences. Bioinformatics 22, 1658–1659 (2006).

50. Katoh, K. & Standley, D. M. MAFFT multiple sequence alignment software version 7: Improvements in performance and usability. Mol. Biol. Evol. 30, 772–780 (2013).

51. Capella-Gutiérrez, S., Silla-Martínez, J. M. & Gabaldón, T. trimAl: A tool for automated alignment trimming in large-scale phylogenetic analyses. Bioinformatics 25, 1972–1973 (2009).

52. Waterhouse, A. M., Procter, J. B., Martin, D. M. A., Clamp, M. & Barton, G. J. Jalview Version 2-A multiple sequence alignment editor and analysis workbench. Bioinformatics 25, 1189–1191 (2009).

53. Quang Minh, B., et al. IQ-TREE 2: New models and efficient methods for phylogenetic inference in the genomic era. Mol. Biol. Evol. 37, 1530–1534 (2020).

54. Kalyaanamoorthy, S., Minh, B. Q., Wong, T. K. F., Von Haeseler, A. & Jermiin, L. S. ModelFinder: Fast model selection for accurate phylogenetic estimates. Nat. Methods 14, 587–589 (2017).

55. Hoang, D. T., Chernomor, O., Von Haeseler, A., Quang Minh, B. & Sy Vinh, L. UFBoot2: Improving the Ultrafast Bootstrap Approximation. Mol. Biol. Evol. 35, 518–522 (2017).

56. Shen, X. X., Li, Y., Hittinger, C. T., Chen, X. xin & Rokas, A. An investigation of irreproducibility in maximum likelihood phylogenetic inference. Nat. Commun. 11, (2020).

57. HMMER 3.3.2 (Nov 2020); http://hmmer.org/ Copyright (C) 2020 Howard Hughes Medical Institute.

58. Parks, D. H. et al. GTDB release 10: a complete and systematic taxonomy for 715 230 bacterial and 17 245 archaeal genomes. Nucleic Acids Res. 54, D743–D754 (2026).

59. Richter, D. J., et al. EukProt: A database of genome-scale predicted proteins across the diversity of eukaryotes. Peer Community Journal 2, (2022).

60. Camacho, C. et al. BLAST+: Architecture and applications. BMC Bioinformatics 10, (2009).

61. Shannon, P. et al. Cytoscape: A software Environment for integrated models of biomolecular interaction networks. Genome Res. 13, 2498–2504 (2003).

62. Kabsch, W. XDS. Acta Crystallogr. D Biol. Crystallogr. 66, 125–132 (2010).

63. Karplus, P. A. & Diederichs, K. Assessing and maximizing data quality in macromolecular crystallography. Curr. Opin. Struct. Biol. 34, 60–68 (2015).

64. McCoy, A. J. et al. Phaser crystallographic software. J. Appl. Crystallogr. 40, 658–674 (2007).

65. Liebschner, D. et al. Macromolecular structure determination using X-rays, neutrons and electrons: Recent developments in Phenix. Acta Crystallogr. D Struct. Biol. 75, 861–877 (2019).

66. Agirre, J. et al. The CCP4 suite: integrative software for macromolecular crystallography. Acta Crystallogr. D Struct. Biol. 79, 449–461 (2023).

67. Williams, C. J. et al. MolProbity: More and better reference data for improved all-atom structure validation. Protein Science 27, 293–315 (2018).

68. Živković, I., Dulic, M., Kozulic, P., Mocibob, M. & Gruic-Sovulj, I. Kinetic characterization of amino acid activation by aminoacyl-tRNA synthetases using radiolabelled γ-[32P]ATP. FEBS Open Bio 15, 580–586 (2025).

69. Zivkovic, I., Moschner, J., Koksch, B. & Gruic-Sovulj, I. Mechanism of discrimination of isoleucyl-tRNA synthetase against nonproteinogenic α-aminobutyrate and its fluorinated analogues. FEBS Journal 287, 800–813 (2020).

