## Supplementary data for "C-terminal evolutionary remodelling of isoleucyl-tRNA synthetases is a prokaryote-specific strategy for tuning aminoacylation rate"

**Supplementary Figure S1.** Validation and performance of IleRS HMMs.

**Supplementary Figure S2.** Maximum-likelihood phylogenies of individual IleRS domains.

**Supplementary Figure S3.** Unbiased electron density maps are used to build and confirm the presence of bound tRNA.

**Supplementary Figure S4.** Architecture of the IleRS2:*Ec*tRNA$^{Ile}$:ATP complex and comparison with the structures of eukaryotic IleRS2 and bacterial IleRS1 enzymes bound to tRNAs.

**Supplementary Figure S5.** Structural rearrangements occur in *Pm*IleRS2 upon tRNA binding.

**Supplementary Figure S6.** Yeast IleRS2 exhibits more stringent anticodon recognition than *S. aureus* IleRS1 and contacts the D-loop by base-specific interactions.

**Supplementary Figure S7.** Truncation of the INS subdomain does not perturb the IleRS2 fold.

**Supplementary Figure S8.** Complementation of mupirocin-inhibited *E. coli* growth by *Pm*IleRS2 variants at various IPTG-induced expression levels.

**Supplementary Table S1.** Summary of data collection and refinement statistics of the determined structure.

**Supplementary Table S2.** Comparison of *Pm*IleRS1 and *Pm*IleRS2 aminoacylation rates toward tRNA$^{Ile}$ GAU and its variants.

**Supplementary Table S3.** Steady-state kinetic parameters for isoleucine activation by wild-type and mutants *Pm*IleRS2.

**Supplementary Table S4.** Primers used to design *Pm*IleRS2 and *Pm*tRNA$^{Ile}$ GAU mutants.

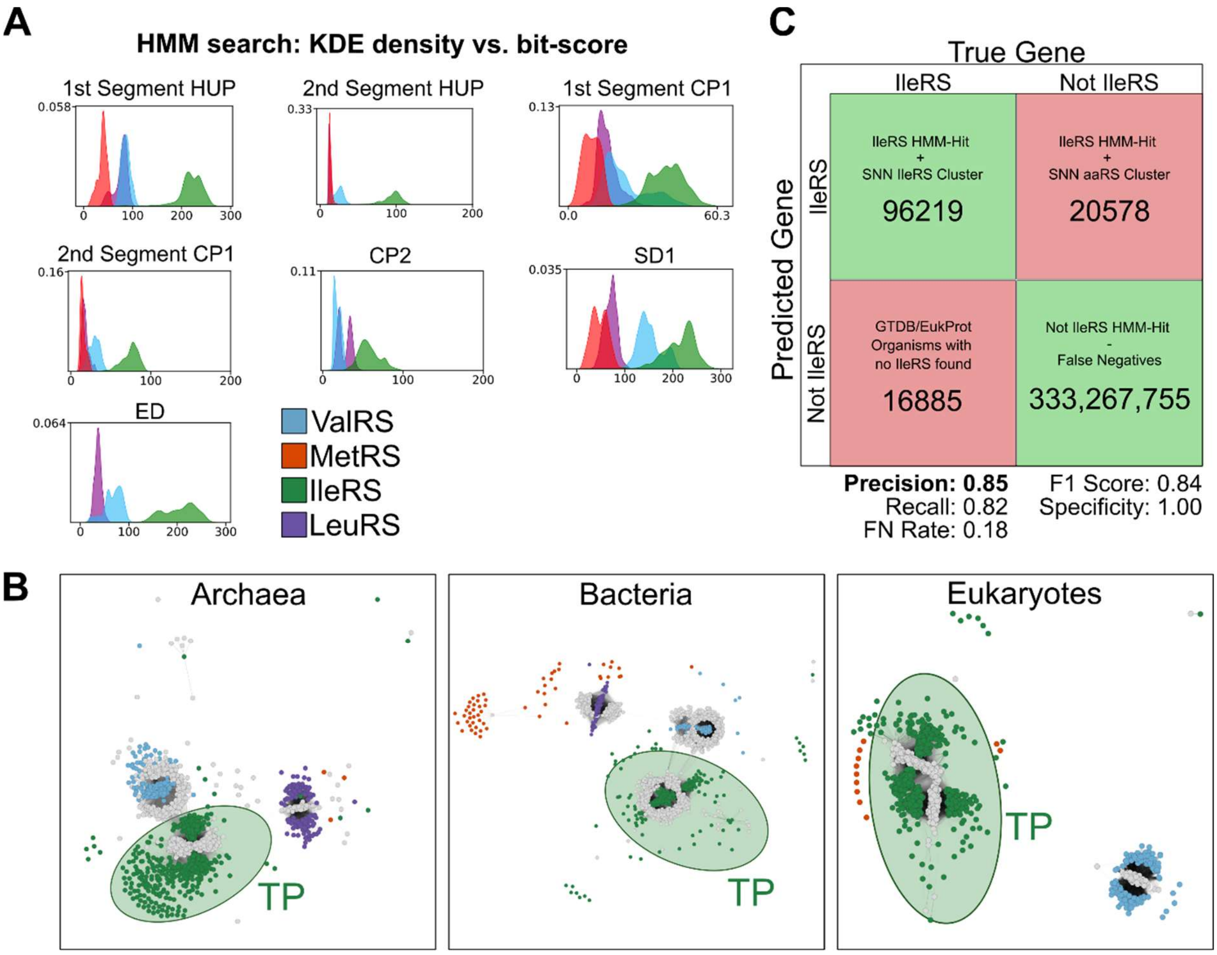


**Figure S1. Validation and performance of IleRS HMMs. (A)** Distribution of normalised domain alignment scores for IleRS, ValRS, LeuRS, and MetRS families, illustrating the discrimination capacity of the HMMs. Each panel corresponds to one IleRS domain HMM, and colour coding reflects the aminoacyl-tRNA synthetase family as indicated in the legend. **(B)** Sequence similarity networks (SSNs) of putative IleRS, validated IleRS, and other class I aminoacyl-tRNA synthetase (AARS) sequences. Nodes represent clustered protein sequences, and edges indicate significant sequence similarity (see Methods). Node colours follow the scheme in (A). Edge shade indicates bit scores; darker edges represent higher similarity and lighter edges represent lower similarity. **(C)** Confusion matrix summarising the classification performance of the IleRS HMMs against the GTDB and EukProt databases. True positives represent sequences clustering with known IleRS genes. False positives correspond to sequences assigned to non-IleRS aminoacyl-tRNA synthetase clusters (ValRS, LeuRS, or MetRS). False negatives denote organisms in which no IleRS genes were detected. True negatives represent all genes searched that did not return an IleRS hit, excluding those estimated as false negatives.

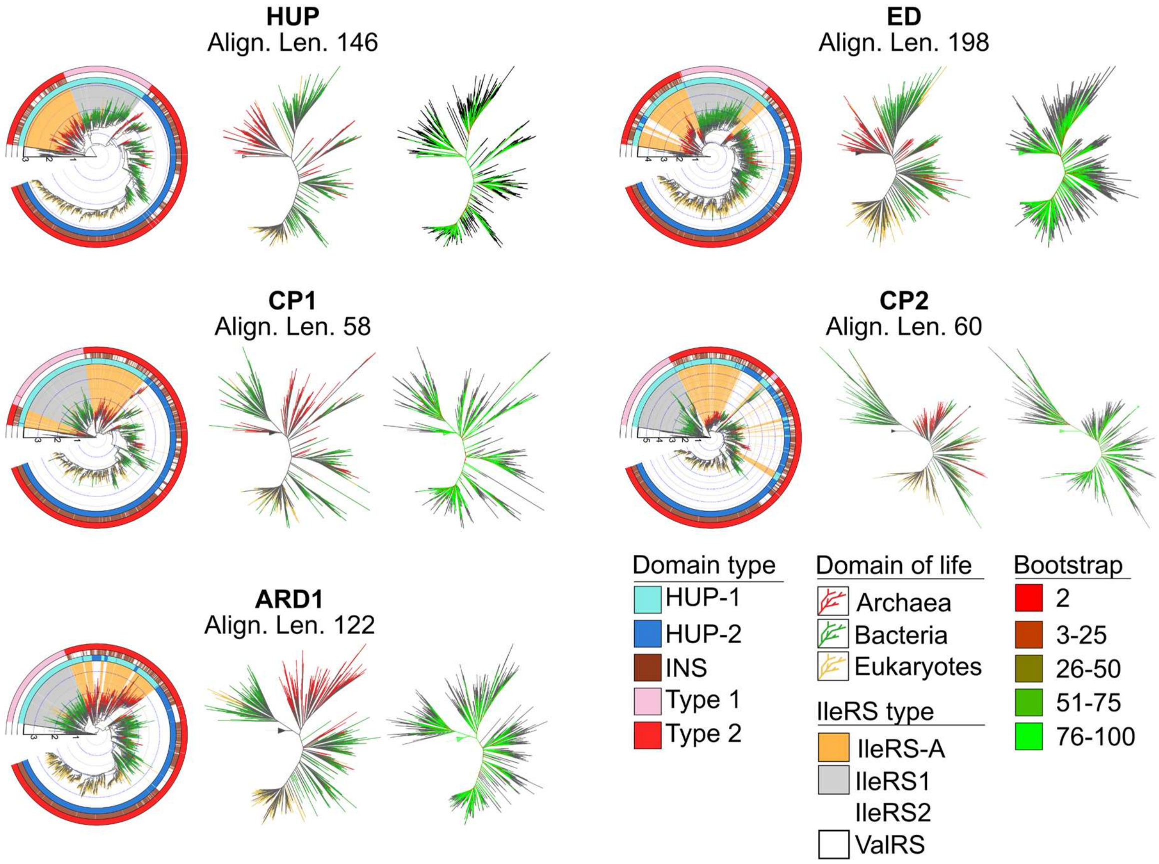


**Figure S2. Maximum-likelihood phylogenies of individual IleRS domains.** Maximum-likelihood trees are shown for the HUP, ED, CP1, CP2, and ARD1 domains, each rooted using the homologous ValRS domain. For each domain, three representations are provided (left to right): a rooted tree with annotated outer rings, an unrooted topology, and an unrooted tree coloured by bootstrap support. In the rooted representations, the concentric outer rings denote (from inner to outer): the HUP domain type, the presence of the INS insertion, and the C-terminal domain type. Branch colours indicate the domain of life (archaea, bacteria, eukaryotes), with bootstrap values displayed using the colour scale shown. Alignment lengths are indicated above each panel. Across the HUP, ED, and CP1 trees, the root consistently falls within archaeal IleRS-A sequences, whereas in CP2 and ARD1, the root is positioned near IleRS-A but between the IleRS1 and IleRS-A clades. In all trees, IleRS-A sequences (characterised by HUP-1 and C-terminal type 2) form a coherent and well-supported cluster, even though neither HUP nor C-terminal sequence information was included in these domain-specific alignments (with the exception of the HUP tree).

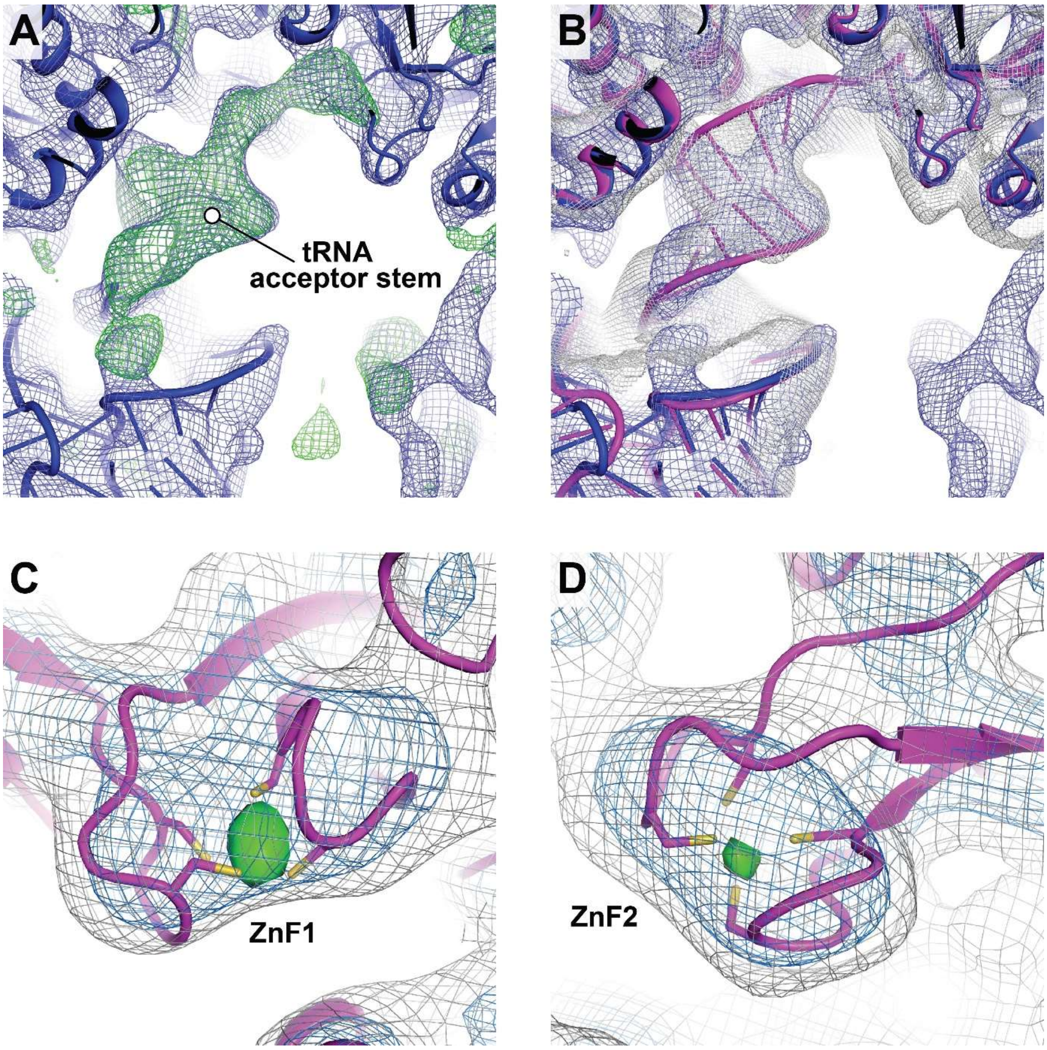


**Figure S3. Unbiased electron density maps are used to build and confirm the presence of bound tRNA: (A)** The initial molecular replacement model (blue cartoon), with parts of the tRNA acceptor stem omitted, showed strong unbiased positive m$F_o$-D$F_c$ Fourier difference density for the missing tRNA segments (green mesh, contoured at 2 $\sigma$) after domain-wise rigid body refinement. The corresponding 2m$F_o$–D$F_c$ Fourier difference map (blue mesh) is contoured at 0.7 $\sigma$. **(B)** Completed model of the acceptor stem of the tRNA molecule. The 2m$F_o$–D$F$c Fourier difference map shown in panel (A) was subjected to 4-fold real-space NCS averaging around the displayed reference molecule, allowing modelling of the missing tRNA residues (magenta) into the averaged map (grey). **(C)** Calculation of the anomalous difference Fourier map (green isosurface, contoured at 4.8 $\sigma$) revealed the positions of the zinc ions in the two zinc fingers (zinc ions are coordinated by C181, C184, C387, and C390 in ZnF1 and C459, C462, C500, and C503 in ZnF2, respectively). The height of the anomalous peaks was used to monitor the progression and convergence of the building and refinement procedure, resulting in considerable improvement of $R_{work}$ and $R_{free}$. Additionally, an unbiased simulated annealing composite OMIT map is shown at 0.8 $\sigma$ and 1.8 $\sigma$ contour levels (grey and blue meshes, respectively).

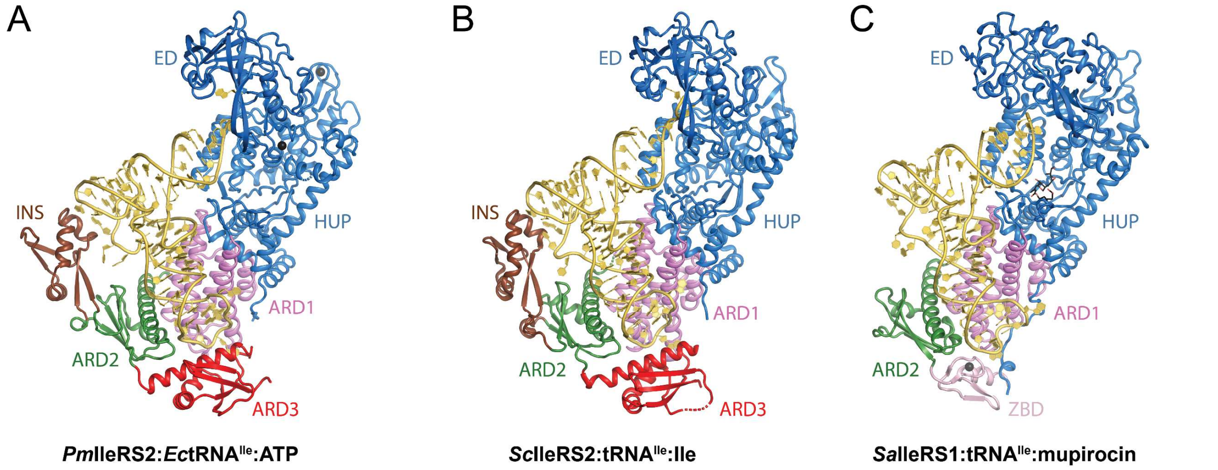


**Figure S4. Architecture of the *Pm*IleRS2:*Ec*tRNA$^{Ile}$:ATP complex and comparison with the structures of eukaryotic IleRS2 and bacterial IleRS1 enzymes bound to tRNAs: (A)** The *Pm*IleRS2:*Ec*tRNA$^{Ile}$:ATP complex features a conserved HUP domain, two connective peptides (CP) hosting the editing domain (ED), and the tRNA anticodon binding domains: ARD1 (violet), ARD2 (green), and ARD3 (red). Insertion into ARD2 (INS, brown) interacts with the tRNA D-loop. The acceptor stem is oriented toward the editing site, while the anticodon loop is positioned within a pocket formed by the ARD1 and ARD3 domains. **(B)** The *S. cerevisiae* IleRS2:tRNA$^{Ile}$:Ile complex (PDB: 8WND) displays an overall architecture analogous to that of *Pm*IleRS2. **(C)** In contrast, the *S. aureus* IleRS1:tRNA$^{Ile}$:mupirocin complex (PDB: 1FFY) lacks INS and features a zinc-binding domain (ZBD, lightpink) at the C-terminus.

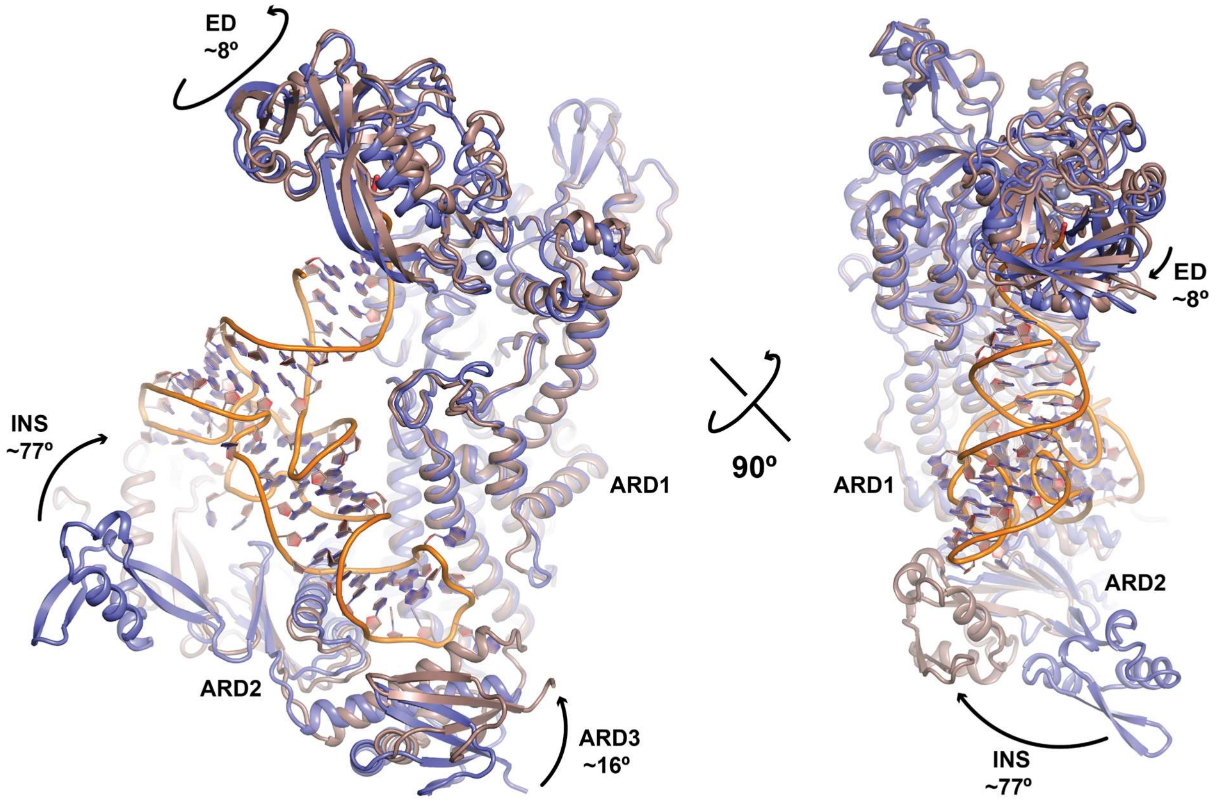


**Figure S5. Structural rearrangements occur in *Pm*IleRS2 upon tRNA binding:** Superposition of the *Pm*IleRS2:*Ec*tRNA$^{Ile}$:ATP complex (brown) with the *Pm*IleRS2:Ile-AMS complex (PDB: 8C8V) (blue), shown in orientations perpendicular to the longest (left) and shortest (right) axes of the complex. The alignment reveals steric rearrangements of *Pm*IleRS2 domains upon tRNA binding. This rearrangement involves repositioning of the editing domain and closure of the subdomains within the C-terminal anticodon-binding domain toward the tRNA. Notably, INS establishes contacts with the D-loop of the tRNA.

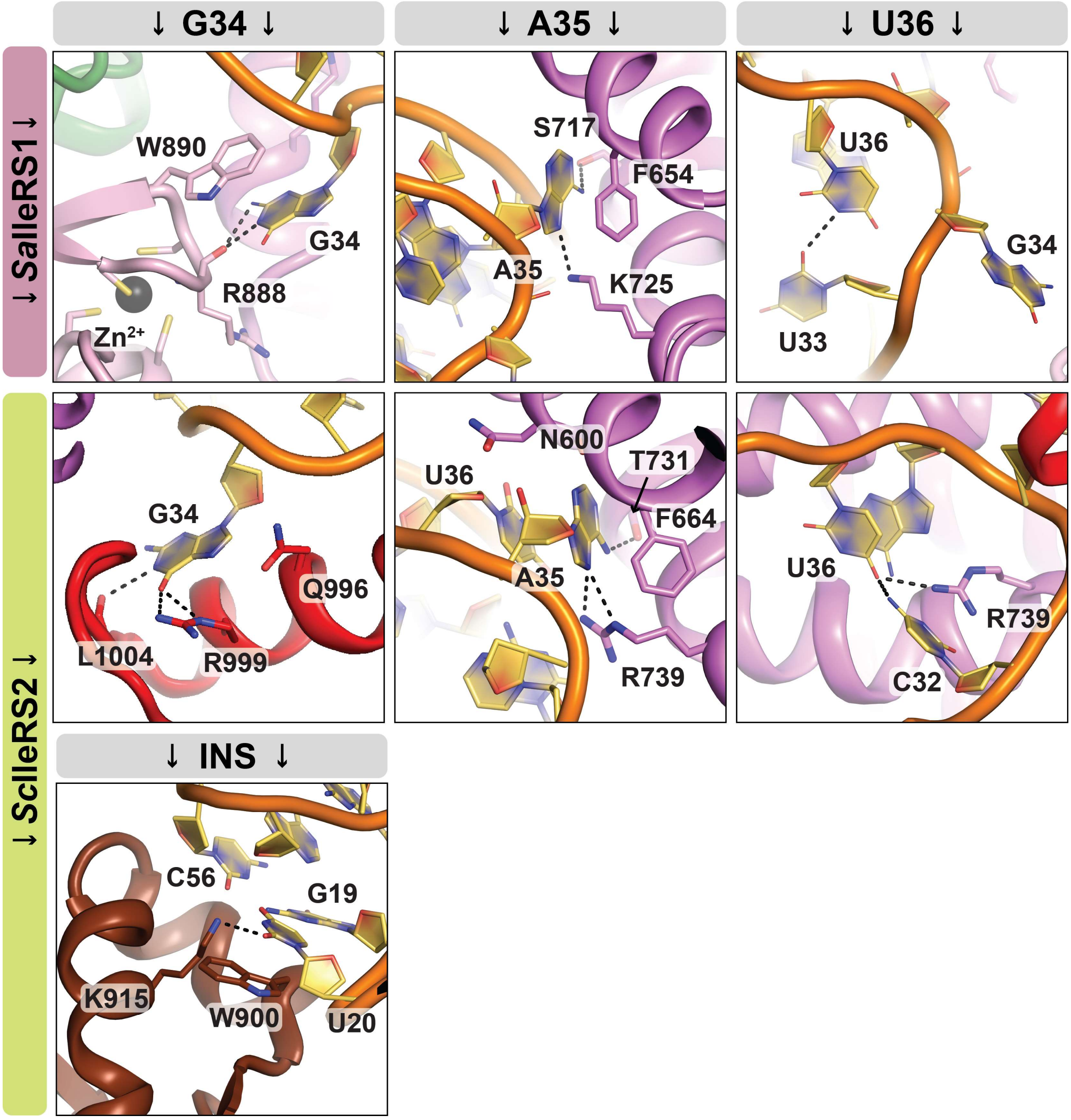


**Figure S6. Yeast IleRS2 exhibits more stringent anticodon recognition than *S. aureus* IleRS1 and contacts the D-loop by base-specific interactions.** Recognition of the three tRNA$^{Ile}$ anticodon bases (G34, A35, U36) by *Sa*IleRS1 (PDB: 1FFY) and *Sc*IleRS2 (PDB: 8WND) is given in the upper and middle panels. The lower panels depict recognition of the tRNA$^{Ile}$ D-loop by *Sc*IleRS2 (PDB: 8WND).

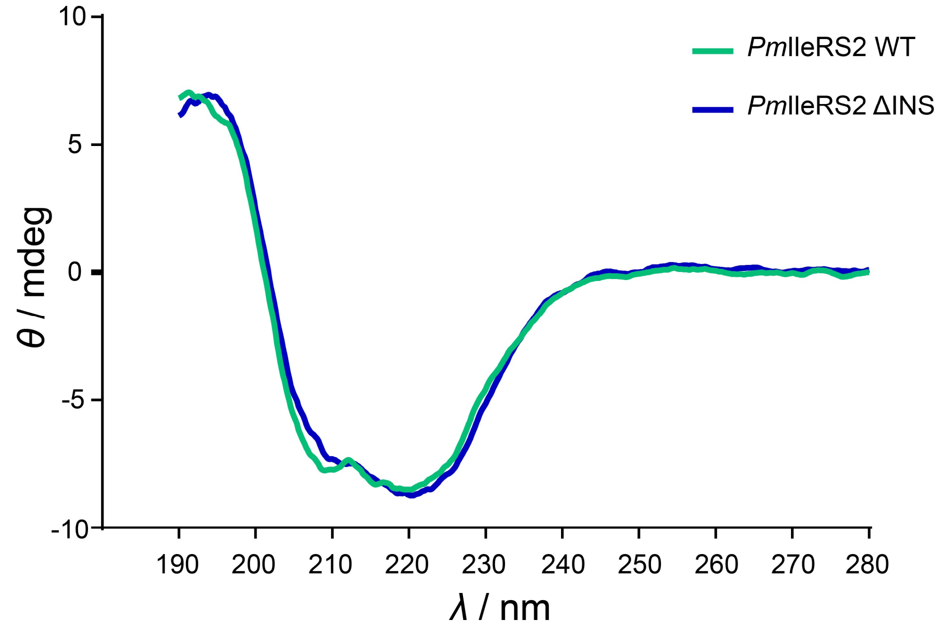


**Figure S7. Truncation of the INS subdomain does not perturb the *Pm*IleRS2 fold.** Circular dichroism (CD) spectrum of *Pm*IleRS2 WT and ΔINS (1 mg mL$^{-1}$) in 1 mM $NaH_2PO_4$ pH = 7.4 at 25 °C; *l* = 0.1 mm; *V* = 30 µL. Spectroscopy of CD did not reveal a structural perturbation related to the excision of INS.

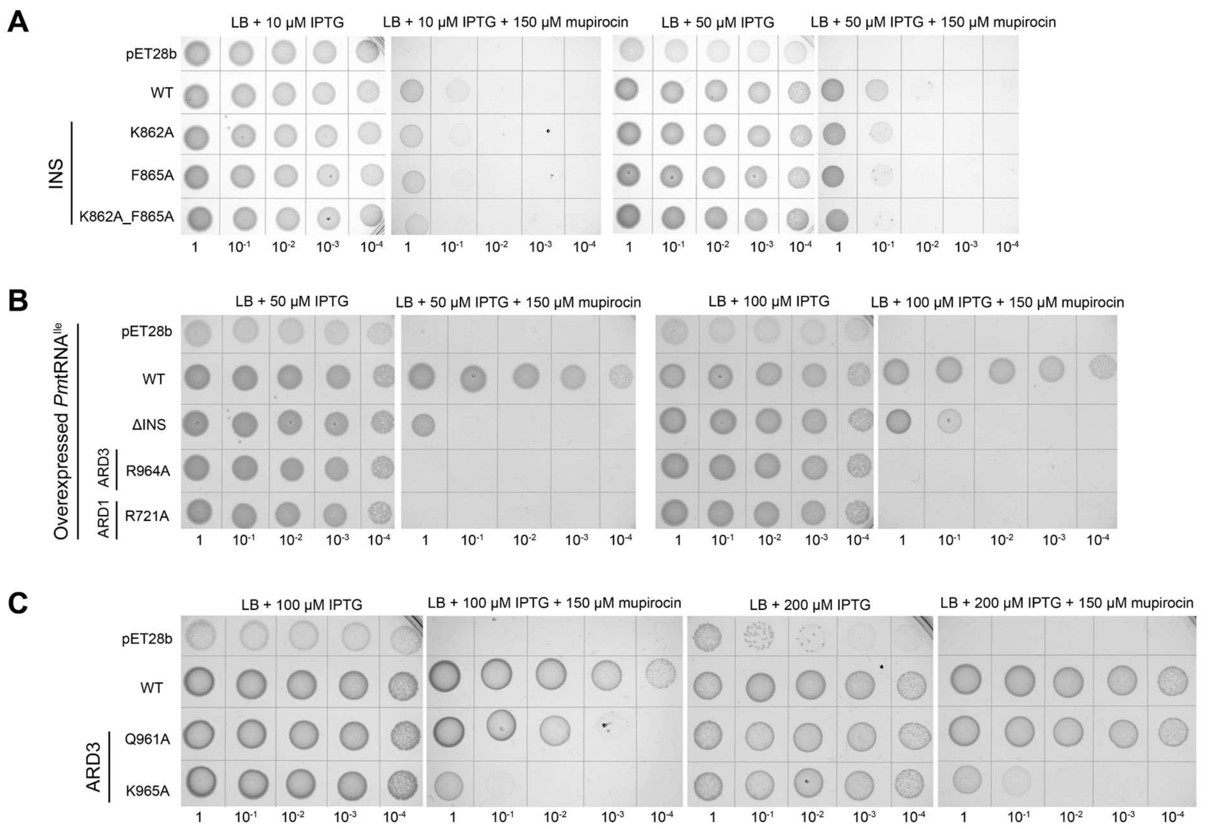


**Figure S8**. **Complementation of mupirocin-inhibited *E. coli* growth by *Pm*IleRS2 variants at various IPTG-induced expression levels.** Functionality of the *Pm*IleRS2 mutants *in vivo* was assessed by their ability to complement mupirocin-inhibited *E. coli* IleRS1. Expression of the mutants was induced from the pET28b plasmid using various IPTG concentrations. **A)** INS mutants rescue *E. coli* growth at all IPTG concentrations used. **B)** ΔINS fail to complement inactivated IleRS1 at 50 μM IPTG, while complementing it weakly at 100 μM IPTG, in both cases, *Pm*tRNA$^{Ile}$ was co-overexpressed. Anticodon-recognising mutants R964A and R721A cannot complement *Ec*IleRS1 in any of these conditions. **C**) Q961A and K965 rescue *E. coli* growth at 100 μM and 200 μM IPTG. For each panel, growth without mupirocin is shown on the left, the negative control (empty plasmid with and without overexpressed tRNA$^{Ile}$) is positioned at the top, and the positive control (overexpression of the WT) is placed in the second row.

**Supplementary Table 1.** Summary of data collection and refinement statistics of the determined structure.

| **Complex** | ***Pm*IleRS2:*Ec*tRNA$^{Ile}$ GAU:ATP** |
|---|---|
| **PDB ID old / extended** | 9U0O / pdb_00009U0O |
| **Data collection and processing** (XDS & XSCALE, BUILD=20220220) | |
| Wavelength (Å) | 1.000 |
| Resolution range* | 48.51-5.30 (5.44-5.30) |
| Space group | $P2_12_12_1$ |
| Unit cell dimensions | a=121.75 Å, b=147.01Å, c=344.11Å, α=β=γ=90° |
| Unique reflections* | 23078 (1660) |
| Multiplicity | 13.5 |
| Completeness (%) | 99.8 (99.6) |
| R-merge (%)* | 10.5 (206.4) |
| **Coordinate refinement** (PHENIX 1.21.2_5419) | |
| $R_{work}$ / $R_{free}$ (%)* | 20.5/25.0 (36.8/44.3) |
| $R_{free}$ test set size (%) | 8.67 |
| Model composition | |
| Non-hydrogen atoms | 39884 |
| Protein residues | 4128 |
| RNA residues | 304 |
| Ligands: $Zn^{2+}$ / $SO_4$ / $Mg^{2+}$ x $6H_2O$ | 8 / 1 / 1 |
| B-factors min/mean/max (Å$^2$) | |
| Protein | 213/364/627 |
| RNA | 257/421/659 |
| Ligands | 247/351/480 |
| **Model Validation** (MolProbity 4.5.2) | |
| *General* | |
| R.m.s. deviations | |
| Bond lengths (Å) | 0.004 |
| Bond angles (°) | 0.72 |
| Clashscore | 5.7 |
| Protein | |
| Rotamer outliers (%) | 0.66 |
| CaBLAM outliers (%) | 2.0 |
| Ramachandran plot | |
| Favored (%) | 95.19 |
| Allowed (%) | 4.54 |
| Disallowed (%) | 0.27 |
| RNA | |
| Pucker outliers (%) | 0 |
| Bond outliers (%) | 0 |
| Angle outliers (%) | 0 |
| Suite outliers (%) | 16.8 |
| (*) Values for highest resolution shell are indicated in parentheses | |

**Supplementary Table 2.** Comparison of *Pm*IleRS1 and *Pm*IleRS2 aminoacylation rates toward $tRNA^{Ile}$ GAU and its variants[a,b].

| | Substrate | $k_{observed}$ / $s^{-1}$ |
|---|---|---|
| IleRS1 | $tRNA^{Ile}$ | 0.41 ± 0.06 |
| | $tRNA^{Ile}$ U20G | 0.29 ± 0.04 |
| | $tRNA^{Ile}$ U21G | 0.30 ± 0.03 |
| | $tRNA^{Ile}$ G19C | 0.31 ± 0.03 |
| | $tRNA^{Ile}$ G34C | 0.026 ± 0.003 |
| | $tRNA^{Ile}$ U36C | 0.025 ± 0.003 |
| | $tRNA^{Ile}$ U36A | 0.029 ± 0.002 |
| | $tRNA^{Ile}$ A35U | 0.010 ± 0.004 |
| IleRS2 | $tRNA^{Ile}$ | 0.33 ± 0.03 |
| | $tRNA^{Ile}$ U20C | 0.47 ± 0.05 |
| | $tRNA^{Ile}$ U21G | 0.58 ± 0.09 |
| | $tRNA^{Ile}$ U21C | 0.60 ± 0.04 |
| | $tRNA^{Ile}$ G19C | 0.46 ± 0.02 |
| | $tRNA^{Ile}$ G34C | 0.0033 ± 0.0003 |
| | $tRNA^{Ile}$ U36C | 0.0015 ± 0.0005 |
| | $tRNA^{Ile}$ U36A | 0.00133 ± 0.00003 |
| | $tRNA^{Ile}$ A35U | 0.0025 ± 0.0005 |

[a]$tRNA^{Ile}$ wild-type and its variants were prepared by *in vitro* transcription.

[b]Observed aminoacylation rate constant under substrate saturation conditions.

Values represent mean ± SEM of at least three independent experiments.

**Supplementary Table 3.** Steady-state kinetic parameters for isoleucine activation by wild-type and mutants *Pm*IleRS2[a].

| | | $k_{cat}$ / $s^{-1}$ | $K_M$ (Ile) / µM | $k_{cat}$ / $K_M$ / $s^{-1}$ $µM^{-1}$ | Relative change[b] $k_{cat}$ / $K_M$ |
|---|---|---|---|---|---|
| | WT | 50.2 ± 1.5 | 64.5 ± 6.7 | 0.78 ± 0.02 | 1 |
| **ARD1** | R721A | 46.6 ± 2.4 | 51.4 ± 9.3 | 0.88 ± 0.04 | 1.1 |
| **ARD3** | R964A | 40.3 ± 2.5 | 58.4 ± 12.4 | 0.7 ± 0.1 | 0.9 |
| | ΔINS | 14.4 ± 0.76 | 61.7 ± 11.3 | 0.24 ± 0.02 | 0.3 |

[a] Activation was tested using the ATP-PPi exchange assay.

[b] Relative change was defined as the ratio of mutants to wild-type enzyme *Pm*IleRS2.

Values represent mean ± SEM of at least three independent experiments.

**Supplementary Table 4.** Primers used to design *Pm*IleRS2 and *Pm*tRNA$^{Ile}$ GAU mutants.

| Primer name | Sequence (5' → 3') |
|---|---|
| ***Pm*IleRS2** | |
| R721A_F | GCCGATCGTTTTTGGTCAGAAGG |
| R721A_R | GGAACGTCGCACATACCAG |
| N647A_F | CTTGTGGCCGTGTACGGCTTTTACG |
| N647A_R | GTACACGGCCACAAGTGTATCAATGAC |
| F651A_F | GTACGGCGCTTACGTGCTGT |
| F651A_R | CGTAAGCGCCGTACACGTTCA |
| S713A_F | AGCTAGCCAACTGGTATGTGCG |
| S713A_R | CCAGTTGGCTAGCTCTTCAATGA |
| Q961A_F | GCAGATTATCGTAAAAAGCTGGATTTACCC |
| Q961A_R | AACCGCTCGAATTACTTCACG |
| K965A_F | GCAAAGCTGGATTTACCCGTGAATTC |
| K965A_R | ACGATAATCTTGAACCGCTCG |
| R964A_F | TCAAGATTATGCTAAAAAGCTGGATTTACCCG |
| R964A_R | ACCGCTCGAATTACTTCAC |
| K862A_F | CGTATTAGCACTGGATTTTAAACAGGCTGG |
| K862A_R | CCAGTGCTAATACGTATGAAACCAGTTTG |
| F865A_F | CTGGATGCTAAACAGGCTGGACCAAAGT |
| F865A_R | GCCTGTTTAGCATCCAGTTTTAATACGTATG |
| K862A_F865A_F | GGTTTCATACGTATTAGCACTGGATGC |
| K862A_F865A_R | GCCTGTTTAGCATCCAGTGCTAATACGT |
| ***Pm*tRNA$^{Ile}$ GAU** | |
| U20G_F | TTATCAGGCGTGCGCTCTACCCAG |
| U20G_R | GCCTATAGCTCAGCTGGGTAGAG |
| U21G_F | TTATCAGGCGTGCGCTCTCACCAG |
| U21G_R | GGCCTATAGCTCAGCTGGTGAGAG |
| G34C_F | ACCGACCTCACGCTTATGAGG |
| G34C_R | GTTAGAGCGCACGCCTCATAAGC |
| A35U_F | ACCGACCTCACGCTTAACAGG |
| A35U_R | GTTAGAGCGCACGCCTGTTAAGC |
| U36A_F | ACCGACCTCACGCTTTTCAGG |
| U36A_R | GTTAGAGCGCACGCCTGAAAAGC |
| U36C_F | ACCGACCTCACGCTTGTCAGG |
| U36C_R | GTTAGAGCGCACGCCTGACAAGC |
| G19C_F | CTAAGCAGCTGAGCTATAG |
| G19C_R | GCTGCTTAGAGCGCAC |

The red nucleotides indicate the mutation sites.
